# GABAergic and glutamatergic synaptic networks and mitochondrial morphology in the thalamic ventral motor and centromedian nuclei of Rhesus Monkey: A comparative 3D Electron Microscopic Analysis between Control and Parkinsonian State

**DOI:** 10.64898/2026.08.20.745566

**Authors:** GJ Masilamoni, R M Villalba, JF Pare, Y Smith

## Abstract

The ventral motor and the centromedian (CM) nuclei receive prominent GABAergic inputs from the basal ganglia, massive glutamatergic projections from motor cortices and significant GABAergic afferents from the reticular thalamic nucleus. There is strong evidence that disrupted processing of information through these connections may contribute to the pathophysiology of the basal ganglia-thalamocortical loop in Parkinson’s disease (PD). To further assess potential ultrastructural changes in synaptic connectivity and mitochondrial integrity that may contribute to these network dysfunctions, we used a 3D electron microscopic approach to determine whether the pattern of synaptic innervation and morphological integrity of dendritic mitochondria are altered in the basal ganglia-receiving parvocellular ventral anterior nucleus (VApc) and CM neurons of MPTP-treated parkinsonian monkeys. Three main conclusions can be drawn from our findings: (1) Although the overall pattern of synaptic innervation of VApc and CM neurons is not altered in parkinsonian monkeys, the volume of putative corticothalamic terminals is significantly increased in both nuclei, (2) the prevalence of corticothalamic terminals in contact with distal dendrites is several orders of magnitude higher in VApc than CM in both control and parkinsonian monkeys, (3) the complexity and ultrastructural integrity of dendritic mitochondria is altered in CM, but not in the VApc, of parkinsonian monkeys. These findings lay the foundation for future studies of changes in cortical neuromodulation of VApc and CM neurons in parkinsonism and suggest that mitochondrial defects may contribute to the degeneration of CM neurons in PD.

**Significance Statement:** Parkinson’s disease is associated with complex functional changes in the neuronal communication between the basal ganglia, thalamus, and cerebral cortex. In this study, Masilamoni et al. used a high-resolution 3D electron microscopy approach to demonstrate morphological changes of corticothalamic terminals in both the ventral motor and centromedian (CM) thalamic nuclei of parkinsonian monkeys. They also provide evidence for decreased complexity and pathology of dendritic mitochondria in CM. These results suggest that morphological changes in the corticothalamic system may contribute to the pathophysiology of the cortico-thalamo-cortical loop in parkinsonism and that mitochondrial pathology may be one of the mechanisms that underlie CM neuronal loss in PD.

## Introduction

Chronic depletion of dopamine from basal ganglia (BG) circuits, as it occurs in Parkinson’s disease (PD), profoundly alters the electrical activity of neurons therein. Disturbed BG outputs have detrimental consequences for their target neurons, especially the motor thalamus (DeLong, 1990; Rubin et al., 2012; Bosch-Bouju et al., 2013). The motor thalamic neurons are key effectors of BG outputs, critically contributing to motor behavior, and their activity is profoundly affected in PD. Recently, we have demonstrated that GPi terminals in the parvocellular ventral anterior nucleus (VApc) and the centromedian nucleus (CM), the two main GPi-recipient motor thalamic nuclei, undergo significant morphometric changes in parkinsonian monkeys (Masilamoni et al., 2024), and others have identified altered electrophysiological activity (including firing rate changes, bursting, oscillatory firing properties, and altered somatosensory responses) in the VApc of animal models of PD and in PD patients (Schneider and Rothblat, 1996; Zirh et al., 1998; Raeva et al., 1999; Magnin et al., 2000; Ni et al., 2000; Guehl et al., 2003; Molnar et al., 2005; Pessiglione et al., 2005; Aymerich et al., 2006; Rolland et al., 2007; Sarnthein and Jeanmonod, 2007; Chen et al., 2010; Bosch-Bouju et al., 2014; Kammermeier et al., 2016). Although the underlying substrate of these electrophysiological changes still remains poorly understood, it is well established that changes in synaptic organization and mitochondrial morphology may play a critical role in regulating cellular metabolism and firing patterns (Chu et al., 2017; Johnson et al., 2017; Iascone et al., 2020; Adoff et al., 2021; Montero et al., 2021; Stoler et al., 2022).

There is strong evidence for synaptic remodeling and pruning of glutamatergic and GABAergic connections throughout the BG-thalamocortical circuitry in animal models of PD (Wichmann and DeLong, 1996; Obeso et al., 2000; Day et al., 2006; Villalba and Smith, 2010; Villalba et al., 2015; Xu et al., 2017; Melief et al., 2018; Villalba and Smith, 2018; Chu, 2020; Swain et al., 2020; Merino-Galan et al., 2022; Ji et al., 2023). In particular, significant changes in the number and ultrastructural features of cortical terminals, associated with robust alterations in electrophysiological and plastic properties of corticostriatal and corticosubthalamic synapses, have been reported in rodent and primate models of PD (Ingham et al., 1998; Day et al., 2006; Raju et al., 2008; Villalba et al., 2009; Villalba and Smith, 2010; Mathai and Smith, 2011; Villalba and Smith, 2011; Mathai et al., 2015; Villalba et al., 2015; Chu et al., 2017; Ji et al., 2023). Results of these studies support the hypothesis that structural and synaptic alterations may contribute to the altered activity of motor thalamic neurons in the parkinsonian state, an issue that remains poorly studied.

Because they are a major source of energy (ATP and NAD+) required for the regulation of Ca^2+^ homoeostasis and signaling, mitochondria are essential components for synaptic transmission, plasticity, and firing rate homeostasis (Duchen, 2000; Nicholls and Budd, 2000; Toescu, 2000; Ruggiero et al., 2021). Mitochondria are transported throughout the neuron to meet the metabolic demands and maintain proper neuronal network functions and are frequently found in axon terminals and dendrites (Delgado et al., 2019). Understanding mitochondrial biology is critical as axons and dendrites are consistently lost before the cell body in neurodegenerative diseases that are thought to involve energy depletion (Li et al., 2001; Scheff et al., 2007; Cheng et al., 2010). Taking into consideration that morphological changes in mitochondria support neuronal plasticity in dendrites and axons (Li et al., 2004; Chang et al., 2006; Rangaraju et al., 2019; Kuzniewska et al., 2020; Wang et al., 2023) combined with the fact that chronically MPTP-treated monkeys undergo CM neuronal loss (Villalba et al., 2014) and that disturbed mitochondrial dynamics are central pathological components of animal models and PD patients (Tanaka et al., 1988; Song et al., 2004; Toomey et al., 2022), another objective of this study is to determine whether dendritic mitochondria undergo pathological changes in chronically MPTP-treated parkinsonian monkeys.

To address these issues, we used high-resolution serial block face/scanning electron microscopy (SBF/SEM) and three-dimensional (3D) reconstruction methods to compare the overall pattern of GABAergic and glutamatergic synaptic innervation of distal and proximal dendrites of thalamocortical neurons in VApc and CM between control and parkinsonian monkeys. Given results from our recent study showing changes in the prevalence of corticothalamic terminals in the VApc and CM of parkinsonian monkeys (Swain et al., 2020), we further assess changes in the corticothalamic system through a detailed ultrastructural analysis of morphological changes of corticothalamic terminals in parkinsonian monkeys. Given the importance of dendritic mitochondria in the regulation of synaptic connectivity and neuronal survival, another objective of this study is to compare the prevalence, complexity and pathology of dendritic mitochondria in the VApc and CM between control and parkinsonian monkeys.

Results of these studies have been presented in abstract forms (Masilamoni et al., 2021).

## Materials and Methods

### Animals

Four adult male rhesus monkeys (Macaca mulatta, 4.5–8.5 kg) from the Emory National Biomedical Research Center colony were used in this study. All procedures were approved by Emory’s Animal Care and Use Committee in accordance with guidelines from the National Institutes of Health. The animals were housed in a temperature-controlled room and exposed to a 12-h light/dark cycle. They were fed twice daily with monkey chow supplemented with fruits or vegetables. The animals had free access to water.

### MPTP administration and evaluation of parkinsonism

The four monkeys used in this study were divided into two groups: two monkeys were drug-naïve, healthy, and served as experimental controls. The other two monkeys were rendered moderately parkinsonian via chronic 1-methyl-4-phenyl-1,2,3,6-tetrahydropyridine (MPTP) regimen as reported elsewhere (Swain et al., 2020; Masilamoni et al., 2024). In summary, the two monkeys were brought into an observation cage and their spontaneous movements within this cage were monitored for 15 min weekly during the MPTP treatment period. Movements within the behavior cage were scored by two experts observers (one of them blinded to the MPTP treatment regimen, monitored, and evaluated from videos) according to a nine-criteria parkinsonism rating scale, with evaluations of gross motor activity, balance, posture, arm bradykinesia, arm hypokinesia, leg bradykinesia, arm hypokinesia, arm tremor, and leg tremor. The differences in the rating scores between the two observers were <6%. The mean values obtained by the two experimenters were used for the study. Each criterion received a score of 0–3 (normal/absent to severe), for a maximal score of 27. Animals were considered stably parkinsonian once they had achieved a score of 10 or higher on the rating scale and a >60% reduction in beam breaks from baseline, when both criteria persisted over 6 weeks following cessation of MPTP treatment. The final rating scores for the two MPTP-treated subjects in this study ranged from 10 to 14, corresponding to moderate parkinsonism (Swain et al., 2020; Masilamoni et al., 2024).

### Intrapallidal AAV5 viral vector injection

To identify GPi terminals projecting to the VApc and CM, the two control and two MPTP-treated monkeys received a total of 2–8µL of AAV5-hSyn-ChR2-EYFP or AAV5-hSyn-Arch3-EYFP in the GPi using surgical procedures described in greater detail in our previous studies (Swain et al., 2020; Masilamoni et al., 2024). Briefly, in the control monkeys, the anterograde viral vector injections were made under isoflurane anesthesia with the animal fixed in a stereotaxic frame using aseptic surgical procedures. Preoperative MRI scans of these monkeys were performed to help define the stereotaxic coordinates to target the sensorimotor GPi territory that projects to VApc and CM. On the other hand, the MPTP-treated parkinsonian monkeys, received viral vector solutions in the GPi using extracellular recordings as a guide to delineate the borders of the neighboring nuclei using procedures previously described from our laboratory (Kliem et al., 2007; Galvan et al., 2010). To optimize the use of nonhuman primates, these two monkeys underwent in vivo electrophysiology recordings of the GPe and GPi neuronal activity before receiving AAV5 injections in the GPi. These two monkeys did not undergo any other procedure or received any drugs besides the collection of the single unit recording data in the normal and parkinsonian state. These animals were euthanized within 1.5–7 months after the viral vector injections (Masilamoni et al., 2024).

### Tissue processing for Serial block face scanning electron microscopy (SBF-SEM)

At the completion of the study, the animals were euthanized with an overdose of pentobarbital, and transcardially perfused with a Ringer’s solution and a mixture of paraformaldehyde (4%) and glutaraldehyde (0.1%). The brains were removed from the skull, postfixed in 4% paraformaldehyde, cut in serial sections (60 µm) with a vibratome and used for post-mortem immunostaining and electron microscopy. The immunostaining and electron microscopy protocols used to localize the Green Fluorescent Protein (GFP) antigens were described in our previous studies (Masilamoni et al., 2024). In brief, thalamic tissue sections containing the GPi-receiving parvocellular ventral anterior (VApc) or the CM nuclei were immunostained with a GFP rabbit antibody (1:5,000 dilution; AB_221569) that also recognizes EYFP to localize anterogradely EYFP-labeled GPi terminals in the VApc and CM. Adjacent sections were immunostained for calbindin D28K antibody (1:4000 dilution; Sigma, catalogue number C9848) to help delineate the location of these nuclei. Selected GFP-immunostained sections went through multi-day staining process for electron microscopy as described earlier, and the sections were put into vials in phosphate buffer solution and sent to the Lerner Research Institute’s 3D EM Core at the Cleveland Clinic (Cleveland, OH) in 4% paraformaldehyde for further SBF-SEM processing.

The tissue samples were then incubated in 1.5% potassium ferrocyanide and 2% osmium tetroxide (in sodium cacodylate buffer, 0.1M) at 4°C, rinsed in double distilled H_2_O, and incubated in 1% thiocarbohydrazide at 60°C, followed by washes in ddH2O. Next, the tissue was placed in 2% osmium tetroxide on a rotator at RT, rinsed in ddH2O, and left at 4°C for 48-72 hours in a saturated aqueous solution of uranyl acetate and then lead stained at 60°C, rinsed in ddH2O, and dehydrated in a series of graded ethanol and propylene oxide. Finally, the tissue was placed in the resin (EMbed-812), first a mixture of resin and propylene oxide (50/50) and after that, in fresh resin (100%) and embedded in Pelco silicone mold at 60°C until the resin was fully cured (8-14 hours). Next, the resin blocks were removed from the mold, trimmed, and mounted on an aluminum pin where the vertical sides of the sample were covered with colloidal silver liquid. Once the imaging surface of the tissue was exposed, the sample was then set up in the SBF-SEM in-chamber ultramicrotome and examined in the electron microscope. Low, medium, and high-resolution 2D images were initially taken of the block face surface to assess the tissue preservation. Multiple series of 200-300 EM images were then obtained using two SBF-SEM systems: a Zeiss Sigma VP scanning EM with Gatan 3View system, and a ThermoFisher Teneo VolumeScope (Pan et al., 2023; Masilamoni et al., 2024).

### 3D Reconstruction from Serial Sectioning Electron Microscopy

Selected areas in the VApc and CM with dense GFP-labeled GPi terminals were used to obtain serial ultrastructural images using an SBF/SEM approach. The location of tissue samples taken for SBF/SEM processing were described detail in our previous paper (Masilamoni et al., 2024). Approximately 200-400 serially scanned micrographic images (∼70 nm-thick) were collected from each region of interest. The postsynaptic targets of terminals were identified as cell bodies, distal, and proximal (<1.0 and >1.0 μm in diameter, respectively) dendrites based on their ultrastructural features and cross sectional diameters (Peters et al., 1991). The Z-trace tool (Reconstruct) was used to measure the diameter of dendrite. From these images, a total of 40 distal (20 control, 20 MPTP) and 12 proximal (6 control, 6 MPTP) dendrites were reconstructed using the 3D software Reconstruct (NIH and synapses.clm.utexas.edu) in VApc, while 42 distal (22 control and 20 MPTP) and 12 proximal (7 control and 5 MPTP) dendrites were reconstructed in CM. Dendrites of projection neurons were differentiated from dendrites of interneurons based on the absence of dendritic vesicles, which are exclusively found in interneurons (Hamori et al., 1974; Sherman and Friedlander, 1988; Jones, 2002; Sherman, 2004; Jones, 2007). Although not analyzed through its full extent, the synaptic innervation of 8 projection neurons’ perikarya, [4 in VApc (3 control and 1 MPTP) and 4 in CM (3 control and 1 MPTP)] was reconstructed. These dendritic profiles and cell bodies were randomly chosen based on the quality of their ultrastructural preservation and extent through which they could be followed and reconstructed through serial images. The criterion was set at a series of images composed of minimum 125, 75, and 40 ultrathin sections for distal, proximal and cell body, respectively.

### 3D reconstructions and quantitative analysis

#### Classification of synapses, identification of the postsynaptic target, and morphological measurements

From the reconstructed dendrites and cell bodies in the VApc and CM, 3 types of axon terminal subtypes were identified based on ultrastructural features reported in previous electron microscopic (EM) studies of the mammalian motor thalamus from our laboratory and others (Ilinsky et al., 1997; Kultas-Ilinsky et al., 1997; Jones, 2002; Swain et al., 2020; Masilamoni et al., 2024): (i) Small terminals with round synaptic vesicles forming asymmetric synapses (RS-type terminals; Figure 1A-D), which largely originate from the cerebral cortex, (ii) terminals with flat/pleomorphic vesicles forming single symmetric synapses (“F1”-type terminals; Figure 1E-H), mainly representing GABAergic terminals from the thalamic reticular nucleus (RTN) or GABAergic interneurons, were considered as putative non-pallidal GABAergic boutons, and (iii) large terminals forming multiple symmetric synapses (“F3”-type terminals; Figure 1I-L), which originate from GABAergic neurons in GPi. These ultrastructural features, combined with the morphometric data collected from the EYFP-labeled pallidothalamic terminals, further confirmed that GPi is the source of both labeled and unlabeled terminals forming multiple symmetric synapses in VApc and CM in control and parkinsonian monkeys. The synapses were characterized as circumscribed membrane specializations with dense material in a wide synaptic cleft between the presynaptic and postsynaptic membranes with aggregates of vesicles at the presynaptic membrane. Synapses with prominent postsynaptic density (PSD) are classified as asymmetric synapses, while those with thin PSDs are classified as symmetric synapses. Since the 3D reconstruct software allows navigation through the stack of images, it was possible to unambiguously identify every single synapse as asymmetric or symmetric based on the thickness of the PSD (Merchan-Perez et al., 2009). Each compartment was colored and sorted into a different list: soma, axons, dendrites, or excitatory or inhibitory pre-synaptic structures. From these serially identified elements, the software created a 3D representation of each object from which it calculated the volume of dendrites and terminals.

**Figure 1.**
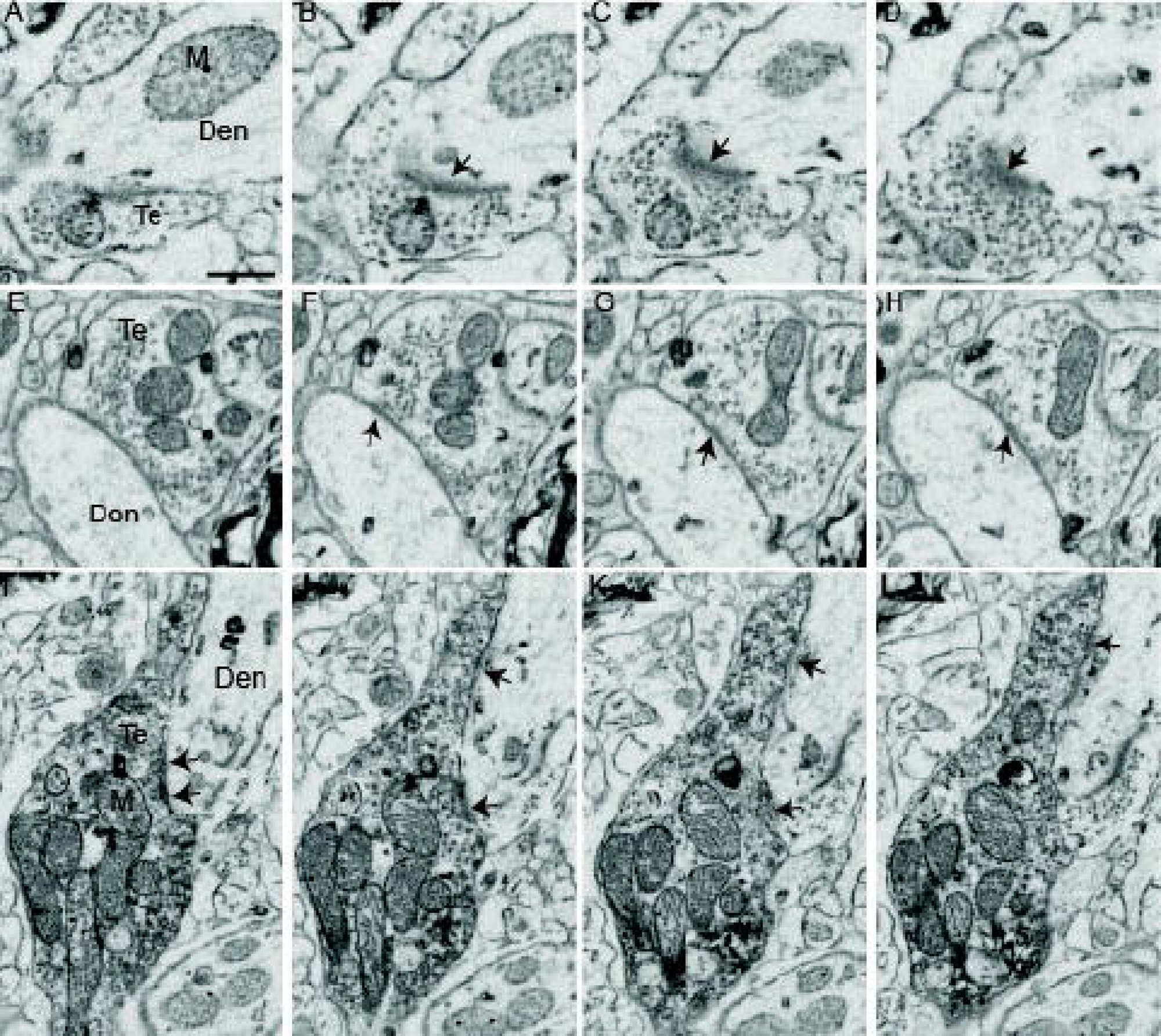
Serial SBF/SEM images of the three main types of axon terminals in the monkey VApc. (A-D) “RS-type” terminals (putative cortico-thalamic) are small-sized, packed with electron lucent vesicles and form asymmetric synapses, (E-H) “F1-type” GABAergic non-pallidal terminals (putatively from RTN or GABAergic interneurons) are medium-sized and form single symmetric synapse and (I-L) “F3-type” large GABAergic terminals from GPi (labeled with AAV5-GFP) that form multiple symmetric synapses (black arrows). Synapse classification was based on examination of the full sequence of serial images (see ref Masilamoni *et al*. (2024) for more detail). Abbreviations: Den: Dendrite; Te: Terminal; Scale bars in A and I are valid for micrographs displayed in A-H and I-L respectively. Scale bars: A and L = 1μm.

#### 3D analysis of dendritic mitochondria

Given the critical role of dendritic mitochondria in normal neuronal function and synaptogenesis and the evidence for CM neuronal loss in PD and chronically MPTP-treated monkeys (Henderson et al., 2000; Villalba et al., 2014), we compared the dendritic mitochondrial morphology in VApc and CM between control and MPTP-treated monkeys. To do so, mitochondria profiles were identified, traced and 3D reconstructed through all stacks of images so that their number and volume could be quantified. Given that individual mitochondria are complex three-dimensional organelles that can be followed across several thin sections, 3D EM reconstruction is essential for rigorously analyzing their morphology and prevalence in brain tissue. For example, in the serially sectioned image stack of dendritic mitochondria shown in Figures 2A-C, mitochondrion M1 appears as a single distinct structure in Figure 2A; however, in subsequent sections, it appears divided into two separate mitochondrial profiles (Fig. 2B-C). Three-dimensional reconstruction confirmed that these apparently separate profiles belong to the same mitochondrion and were therefore counted as a single mitochondrion in our analysis (Fig. 2D).

**Figure 2.**
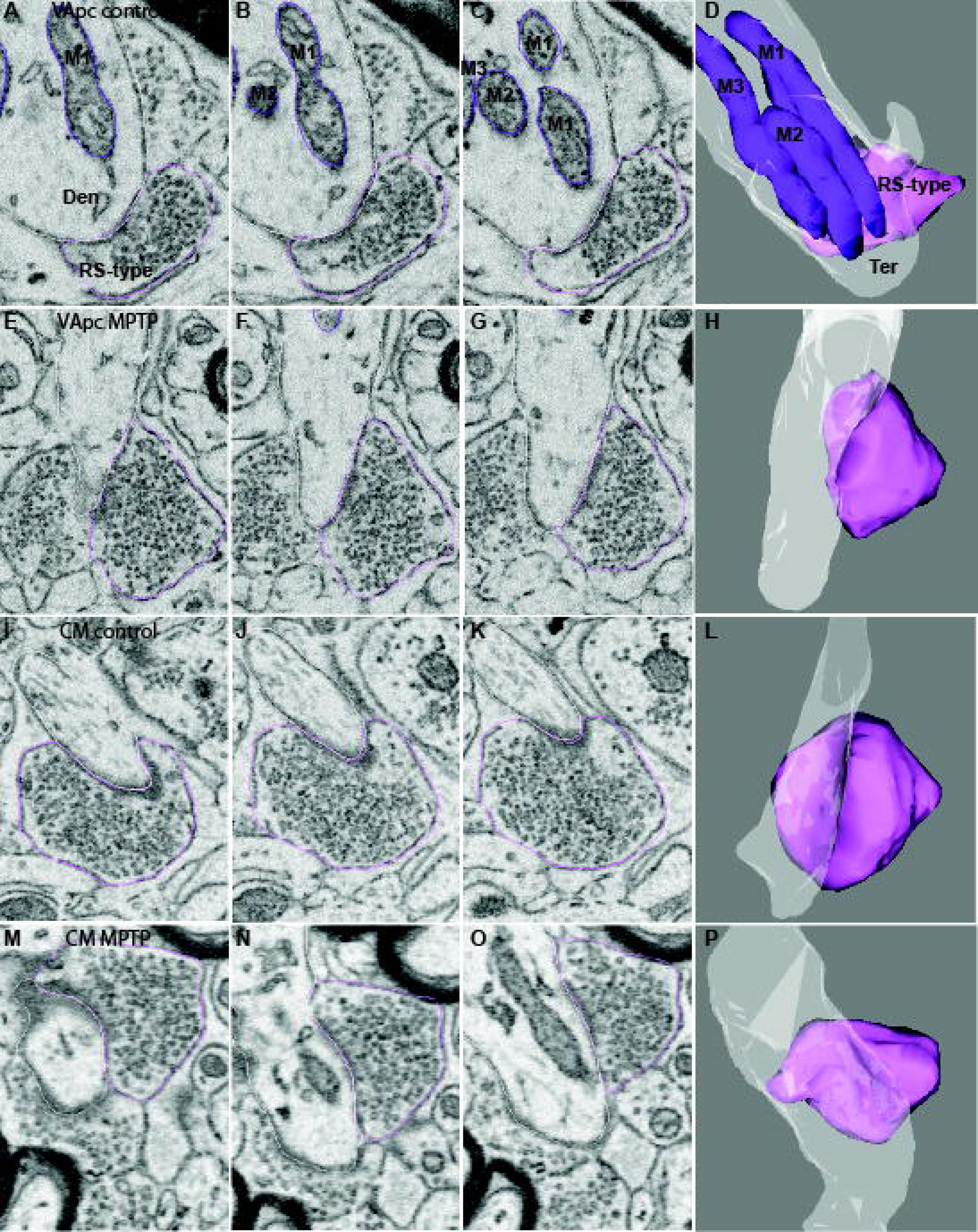
Serial SBF/SEM images and corresponding 3D reconstructions (A-P) of cortical (RS-type) axon terminals forming asymmetric axo-dendritic synapses in the ventral anterior parvocellular (VApc) and centromedian (CM) thalamic nuclei of control and MPTP-treated monkeys. For the axo-dendritic complex shown in panels A-D, dendritic mitochondria have also been reconstructed to illustrate the complex tri-dimensional structure of mitochondria profiles and highlight the need of 3D EM approach to morphometrically study and numerically quantify these organelles. For instance, when examined in panel A it appears that the dendritic segment includes a single mitochondrion, but in subsequent images B and C, the mitochondrion begins to divide and forms two distinct mitochondria. Taking advantage of the SBF/SEM approach, we traced individual mitochondria across serial sections to generate 3D reconstructions (D) from which quantitative analysis and comparison of mitochondrial morphometry between control and MPTP-treated parkinsonian monkey can be performed. In the 3D reconstructions, dendrites are shown in white, RS-type terminals in pink, and mitochondria in purple.

We used the mitochondrial complexity index (MCI) to quantify changes in the mitochondrial shape complexity (Vincent et al., 2019; Faitg et al., 2021). The MCI was calculated using the following formula MCI = SA3/16π2V2 where SA is the surface area and V is the volume of the mitochondria. This equation is used to assess the mitochondrial morphological complexity irrespective of volume (Vincent et al., 2019), thereby providing a quantitative parameter to characterize dissimilarities in mitochondrial morphology between control and parkinsonian states.

#### Statistical Analysis

The data were statistically analyzed using the GraphPad Prism software (version 10.3). Because of the small number of monkeys (two control and two MPTP-treated) used in this study, individual cellular elements (instead of animals) were used in the comparative statistical analyses of changes in terminals density in contact with dendrites and cell bodies of thalamocortical neurons and dendritic mitochondria volume in VApc and CM between control and MPTP-treated parkinsonian monkeys. To reduce the likelihood that the data from one animal drove statistical group differences, unpaired t tests were achieved to determine variability between mean values and variances from animals in the same group. This analysis revealed no significant differences between the data from the two control and MPTP-treated monkeys for both VApc and CM (Table 1). Given these results, data from both animals in each experimental group were pooled for analysis (Masilamoni et al., 2024). Multiple comparisons for two-way ANOVA for repeated measures followed by Sidak’s post hoc test were used to compare the relative abundance of cortical (RS), putative reticular (F1) and pallidal GABAergic (F3) terminals, and mitochondria volume per terminal, between control and MPTP-treated animals. Significance was taken at p<0.05*, p < 0.001**, and p < 0.0001***. All results are expressed as mean ± standard deviation (SD). Cumulative frequency distribution of cortical terminals and dendritic mitochondrial volumes were plotted in a violin plot, and unpaired t tests were performed to determine variability between mean values and variances between control and MPTP-treated monkeys.

## Results

### Nigrostriatal dopamine denervation in MPTP-treated monkeys

In the present study we used chronic low dose of MPTP exposure to slowly induce progressive parkinsonian motor signs and nigrostriatal dopaminergic denervation in two monkeys. A detailed description of this MPTP treatment protocol, quantitative data about parkinsonian motor scores, extent of striatal dopamine denervation, and nigral dopaminergic neurons loss are provided in our previous studies (Swain et al., 2020; Masilamoni et al., 2024). In brief, after the last injection of MPTP, the monkeys displayed stable moderate parkinsonian symptoms, including akinesia, bradykinesia, postural instability for at least 6 weeks prior to sacrifice. In postmortem tissue, TH immunostaining was significantly decreased in the whole striatum, but most particularly in the post-commissural and anterolateral putamen. Albeit some denervation, the caudate nucleus, anteromedial putamen and nucleus accumbens were less affected (Suppl. Fig. 1). As expected, at the midbrain level, the ventral tier of the substantia nigra pars compacta (SNc) were severely damaged, whereas a significant number of TH-immunoreactive neurons and processes remained in the ventral tegmental area and dorsal tier SNc (Swain et al., 2020; Masilamoni et al., 2024) (Suppl. Fig. 1E-F).

### Synaptic inputs to dendrites and cell body of VApc and CM neurons in control and parkinsonian monkeys

Our sample preparation and imaging protocol produced SBF-SEM images with high structural integrity from which the membrane contours of dendrites, and presynaptic axons were clearly visible. Synaptic contacts and post-synaptic density (PSD) of asymmetric excitatory synapses were clearly discernible in VApc and CM of control and MPTP-treated monkeys (Figure. 1A-D). As mentioned above, RS terminals were small- to medium-sized, packed with round synaptic vesicles and occasional mitochondria and formed asymmetric synapses (Figure 1A-D). On the other hand, pallidothalamic terminals (called F3 terminals), which were identified based on specific ultrastructural features reported in previous studies (Ilinsky et al., 1997; Kultas-Ilinsky et al., 1997; Sidibe et al., 1997) and their anterograde labeling from GPi (Swain et al, 2020), were large (1.0–3.0 μm in diameter), enriched in mitochondria, and formed multiple synapses predominantly with dendritic profiles devoid of synaptic vesicles (i.e., of projection neurons) in VApc and CM (Fig. 1 I-L). On the other hand, the non-pallidal, putative reticular GABAergic terminals (called F1 terminals) forming single symmetric synapses (Ilinsky et al., 1999), were much smaller in size and displayed lower mitochondrial volume than GPi terminals in VApc and CM (Fig. 1E-H).

Using these well-established ultrastructural criteria, we compared the density of the different population of terminals in contact with distal dendrites, proximal dendrites and cell bodies of VApc and CM projection neurons between control and parkinsonian monkeys. To do so, we reconstructed the total synaptic innervation of 106 dendritic profiles in VApc and CM of control and parkinsonian monkeys. The average length (in μm) of reconstructed distal and proximal dendrites were 10.33 ± 2.94 and 6.3 ± 1.99 in the VApc of control monkeys, 9.84 ± 1.50 and 5.08 ± 2.37 in the VApc of parkinsonian monkeys; 10.71 ± 2.98 and 7.0 ± 1.86 in the CM of control monkeys and 10.0 ± 1.52 and 8.66 ± 1.14 the CM of parkinsonian monkeys. Table 2 provides additional details about the total number and length of dendritic profiles and number of afferent terminals examined in both groups of monkeys.

**Table 2:** Number of dendrites, cell bodies, terminals, and mitochondria reconstructed and analyzed in this study.

| Group | Thalamic nuclei | Subcellular compartments |  | Types of terminals |  |  | Dendritic mitochondria |
| --- | --- | --- | --- | --- | --- | --- | --- |
|  |  |  |  | RS | F1 | F3 |  |
| Control | VApc | Distal | 20 | 610 | 103 | 13 | 107 |
|  |  | Proximal | 6 | 14 | 8 | 12 | 140 |
|  |  | Cell body | 3 | 13 | 4 | 6 | NA |
|  | CM | Distal | 22 | 188 | 136 | 32 | 142 |
|  |  | Proximal | 7 | 18 | 34 | 16 | 147 |
|  |  | Cell body | 3 | 16 | 27 | 13 | NA |
| MPTP | VApc | Distal | 20 | 600 | 68 | 17 | 92 |
|  |  | Proximal | 6 | 10 | 3 | 9 | 36 |
|  |  | Cell body | 1 | 5 | 1 | 1 | NA |
|  | CM | Distal | 20 | 137 | 76 | 11 | 193 |
|  |  | Proximal | 5 | 15 | 26 | 13 | 210 |
|  |  | Cell body | 1 | 2 | 1 | 3 | NA |

As described in previous studies (Ilinsky et al., 1997; Kultas-Ilinsky et al., 1997; Sidibe et al., 1997; Jones 2007), each dendritic profile received inputs from a variable number and density of RS, F1, and F3 terminals in control and MPTP-treated monkeys. Figure 3A-H shows representative 3D reconstructed distal and proximal dendrites from VApc and CM of control and MPTP-treated parkinsonian monkey. Overall, the density of RS terminals was significantly higher than that of F1 and F3 terminals on the distal dendrites of both VApc and CM neurons in control and parkinsonian monkeys, whereas F1 and F3 terminals were far less abundant and more evenly distributed along the whole somatodendritic domain of these neurons (Figs 3A-H, 4A-B). Furthermore, the present data demonstrate that the density of RS-, F1-, and F3-type terminals on the cell body is considerably lower than that observed on the dendrites (Suppl. Fig.2). Two important findings came out of this analysis: (1) The density of RS terminals in contact with distal dendrites of VApc neurons is significantly higher than the density of RS inputs to distal dendrites of CM neurons (Fig. 4C,D and Suppl. Fig.2), (2) No significant group difference was found for RS, F1 and F3 terminal densities between control and parkinsonian monkeys in both thalamic nuclei (Figs 4A,B; Table 1).

**Figure. 3.**
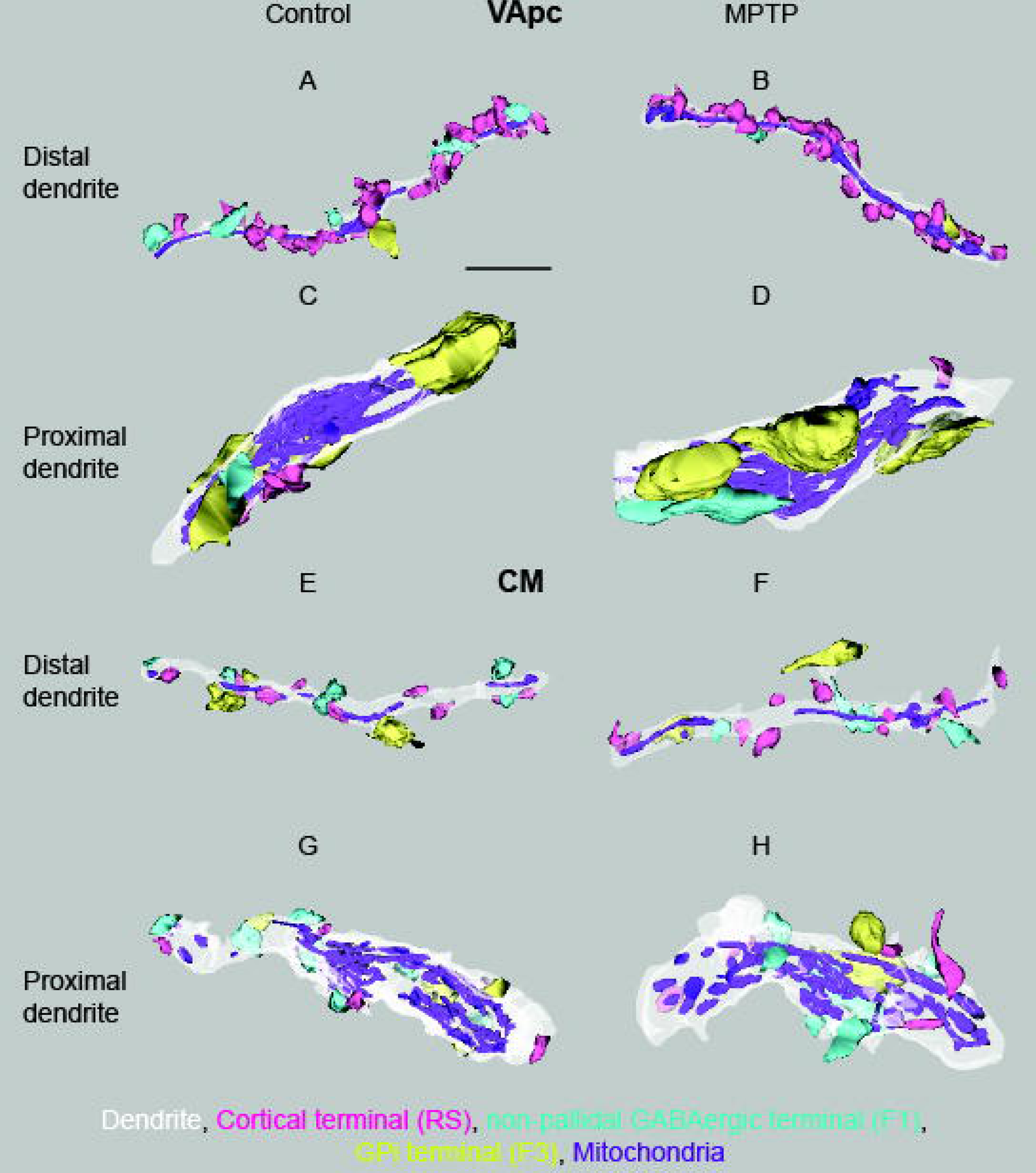
3D reconstruction of the relative prevalence and distribution of RS, F1 and F3 terminals on distal and proximal dendrites of VApc and CM neurons in control and parkinsonian monkeys (A-H). Scale bars: A–H = 2μm. In the 3D reconstructions, dendrites are shown in white, RS-type terminals in pink, F1-type terminals in turquoise, F3-type terminals in yellow, and mitochondria in purple.

**Figure. 4.**
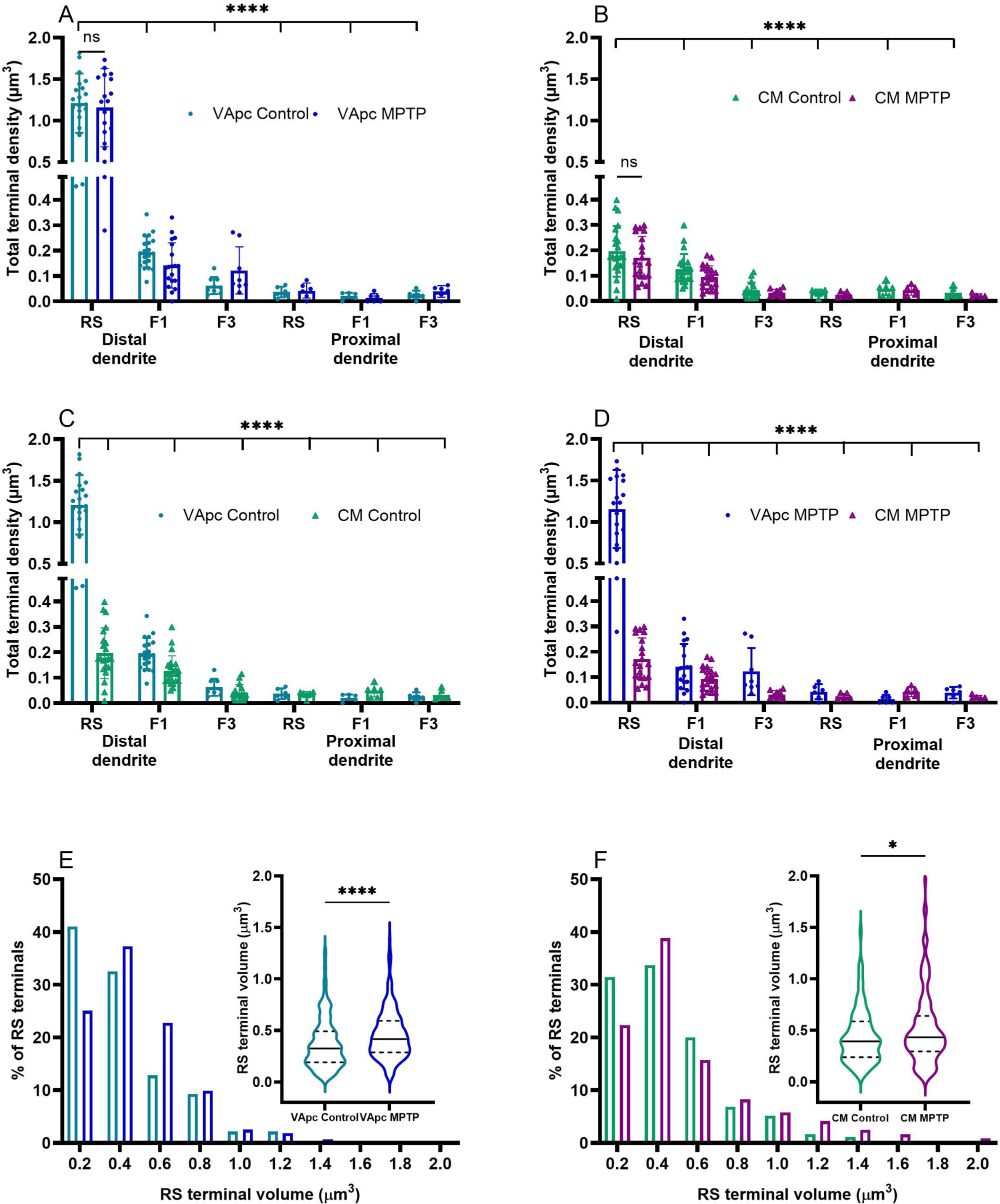
Scatter dot plots with bar graphs comparing the total terminal densities of RS, F1, and F3 terminals in the distal and proximal dendrites of the VApc and CM between control (teal and green) and MPTP-treated parkinsonian monkeys (blue and purple) (A, B). Each data point indicates the density of terminals in contact with individual dendritic profiles (VApc: n = 21 and CM: n=20). No total terminal density differences were found for the three types of terminals between control and parkinsonian monkeys. The density of RS terminals is significantly larger than the density of F1 and F3 terminals in both VApc and CM of control and parkinsonian monkeys (P < 0.0001; A and B). The density of RS terminals in contact with distal dendrites in VApc is significantly larger than in CM (P<0.0001; C and D). E, F, Histograms comparing the relative frequency distribution of RS terminal volumes in VApc (E) and CM (F) between control [n = 275 (VApc, 173, and CM= 112)] and parkinsonian [n = 398 (VApc, 220, and CM: 178)] monkeys. Frequency distribution and violin plot analyses (insert E and F) of RS terminal volumes revealed a significant shift toward larger RS terminals in parkinsonian (MPTP-treated) monkeys compared with controls in both the VApc and CM nuclei. Unpaired t-test analysis confirmed a significant increase in the proportion of large-sized RS terminals following MPTP treatment.

### Morphometric changes of corticothalamic terminals in parkinsonian monkeys

In a recent transmission EM study, we showed a decreased prevalence and an increased diameter of corticothalamic terminals in the VApc and CM of parkinsonian monkeys (Swain et al. 2020). However, the cross-sectional diameter measurements of RS terminals presented in this study were collected from measurements gathered in single ultrathin sections. To further address this issue, we performed an extensive analysis of potential ultrastructural changes of RS terminals (putative corticothalamic) between control and parkinsonian monkeys using a 3D EM reconstruction approach from our SBF/SEM material. Only terminals that could be seen through their full extent in serial sections were reconstructed and analyzed for volume analysis. We reconstructed 741 RS terminals in control monkeys (VApc: 558, CM: 183) and 578 terminals in parkinsonian animals (VApc: 450, CM: 128). Representative serial SBF/SEM images and the corresponding 3D reconstructions of RS terminals are shown in Figure 2A-P. The frequency distribution analysis revealed that the volume of RS terminals varied between 0.2 to 1.2/1.4 μm^3^ for VApc/CM in control and 0.2 to 1.4/1.6 μm^3^ for VApc/CM in parkinsonian monkeys (Fig. 4E-F). We then plotted the RS terminal volume clusters as violin plots (inserts: Fig. 4E-F) and found a significant increase in the average volume of RS terminals in the VApc and CM of MPTP-treated monkeys relative to controls. On average, RS terminal volumes in VApc and CM were 124% and 118% larger in parkinsonian monkeys, respectively (mean volume = 0.46 µm^3^ and 0.52 µm^3^), than in controls (mean volume = 0.37 µm^3^ and 0.44 µm^3^).

### CM dendritic mitochondrial morphology is significantly altered in parkinsonian monkeys

Given evidence from different brain regions that the density and distribution of dendritic mitochondria regulates dendritic morphology, spinogenesis, and the plasticity of spines and synapses (Li et al., 2004) and that alterations in mitochondrial size, shape and number contribute to the pathophysiology of brain disorders (Trimmer et al., 2000; Brustovetsky et al., 2021; Liu et al., 2021; Toomey et al., 2022), we used the 3D SBF/SEM approach to compare dendritic mitochondrial morphology in VApc and CM between control and parkinsonian monkeys. All mitochondria from 106 (55 control and 51 parkinsonian) dendritic profiles (both distal, and proximal) analyzed in this study were fully reconstructed and morphometrically analyzed (Fig 2A-D). A total of 546 (254/292 in control VApc/CM) and 501 (120/381 in parkinsonian VApc/CM) mitochondria were manually traced from each image stack to generate 3D reconstructions. The dendritic mitochondrial volume frequency distribution revealed ∼17% increase in the relative percentage of small-sized (volume ranging from 0.1 to 0.39µm^3^) mitochondria in the CM of parkinsonian monkeys compared to controls, but no change in VApc (Fig 5A-B). We then plotted the frequency components of all dendritic mitochondrial volume clusters as violin plots and found a significant decrease in mitochondrial volume in CM dendrites of parkinsonian monkeys compared with controls (Figure 5B insert), while no significant change was found in VApc (Fig 5A and insert).

**Figure 5.**
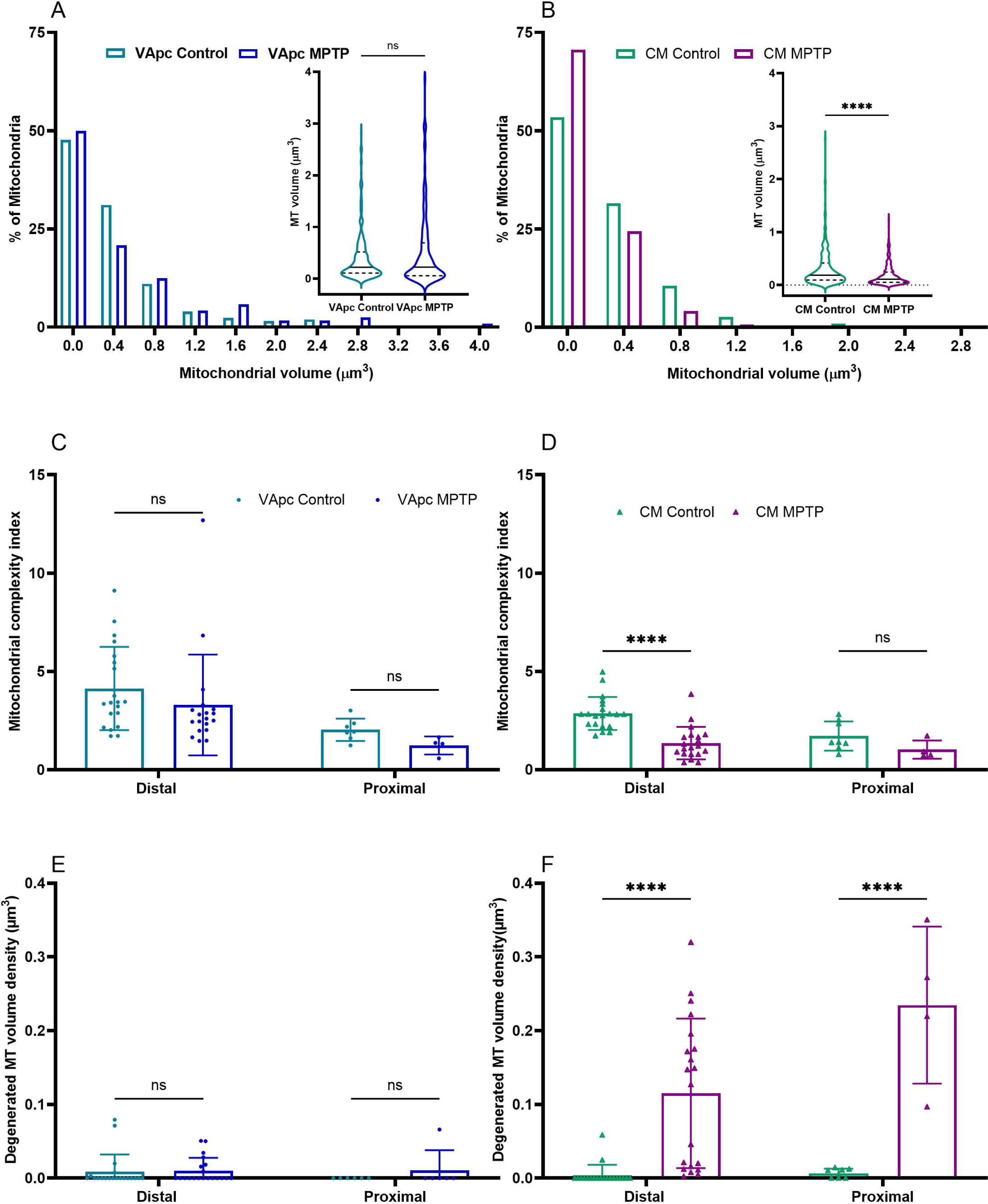
Histograms comparing the relative frequency distribution of mitochondrial volume in the dendrites of VApc (A) and CM (B) neurons between control [n= 546 (VApc, 254, and CM= 292) mitochondria] and parkinsonian [n = 501 (VApc, 120, and CM: 381) mitochondria] animals. Note the higher proportion of smaller mitochondria in CM of parkinsonian monkeys. Violin plots represent the dendritic mitochondrial volume frequency distribution of VApc (insert A) and CM (insert B) between control and parkinsonian monkeys. Unpaired t-test analysis showed a significant increase in the proportion of small-sized mitochondria in CM dendrites, but not in VApc dendrites, following MPTP treatment. C, D. Scatter dot plots with bar graphs comparing the mitochondrial complexity index (MCI) of 3D-reconstructed dendritic mitochondria in the VApc (C) and CM (D) of control and parkinsonian monkeys. The data shows a significant reduction in MCI only in distal CM dendrites following MPTP treatment, whereas no significant changes were observed in proximal CM dendrites or in VApc dendrites. Scatter dot plots with bar graphs comparing the degenerated mitochondrial volume density (DMVD) in 3D-reconstructed dendrites of the VApc (E) and CM (F) of control and parkinsonian monkeys. DMVD was significantly increased in both distal and proximal CM dendrites of parkinsonian monkeys compared with controls (two-way ANOVA with Sidak’s post hoc test, p < 0.0001). In contrast, no significant differences in DMVD were observed between control and parkinsonian monkeys in the VApc.

To further assess potential changes in mitochondrial morphology of dendrite between control and parkinsonian monkeys, we measured MCI (Vincent et al., 2019; Faitg et al., 2021). Based on this metric, mitochondria in the distal dendrites of CM neurons of parkinsonian monkeys were significantly less complex than those of control animals (mean MCI = 2.855, and 1.346, Fig 5D, Table 1). However, no significant MCI change was found in proximal CM dendrites and in both distal and proximal dendritic profiles in VApc (Fig 5C-D, Table 1).

### Mitochondrial electron dense inclusion bodies in CM of parkinsonian monkeys

Mitochondrial abnormalities and intramitochondrial inclusions in neurons of the substantia nigra have been reported in several animal species (monkey, dog, mouse) after MPTP exposure (Tanaka et al., 1988; Christie-Pope et al., 1989; Mizukawa et al., 1990; Rapisardi et al., 1990; Hayes et al., 2021). Given evidence of robust neuronal loss in CM/Pf of PD patients and chronically MPTP-treated monkeys (Henderson et al., 2000; Villalba et al., 2014), we hypothesized that MPTP might induce ultrastructural mitochondrial changes that may disrupt the energy supply and eventually lead to the death of CM/Pf neurons in MPTP-treated monkeys. To address this issue, we reconstructed individual mitochondria through at least 5 serial sections in 20 distal and 5 proximal dendrites in the CM of parkinsonian monkeys and 22 distal and 7 proximal dendrites in controls. As shown in Figure 6, several mitochondria in CM dendrites contained electron dense inclusion bodies. In some cases, there was condensation of mitochondrial content into dense circular structures suggesting early stages of mitochondrial inclusion (white arrowhead, Fig 6A-I). The loss of distinct cristae was also frequently noticed in dendritic mitochondria of MPTP-treated monkeys (black single or double arrowhead, Fig 6A-I). A total of 12 out of 20 distal dendrites and 5 out 6 proximal dendrites in the CM of parkinsonian monkeys exhibited mitochondrial ultrastructural changes, suggesting that these neurons were impaired in the clearance of damaged mitochondria under cellular stress or metabolically compromised. In contrast to CM, the dendritic mitochondria examined in the VApc of control and parkinsonian monkeys did not exhibit significant ultrastructural anomalies (Fig 5C,E), suggesting that MPTP toxicity affects preferentially mitochondrial function in CM neurons of parkinsonian monkeys (Fig 5D,F and 6).

**Figure 6.**
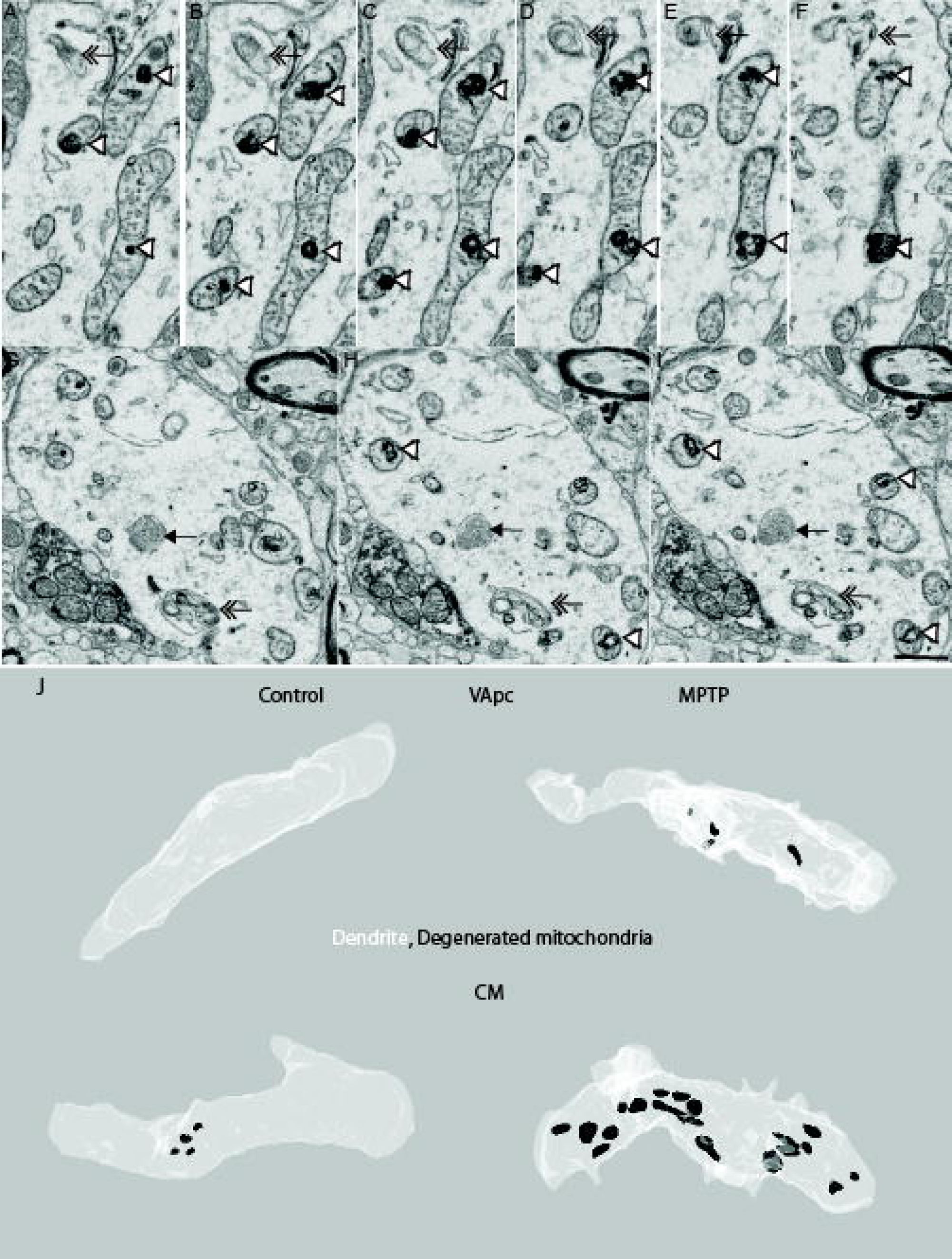
Serial FIB/SEM images of dendritic mitochondria contain electron dense inclusion (white arrowhead), deserted cristae (black arrow) degenerated mitochondria (double arrow): sign of degeneration formed in the CM of a parkinsonian monkey. The classification was based on the examination of the full sequence of serial images (A-F and G-I sequence of serial images). Scale bars: A–H, 1 μm. The reconstructed 3D models of mitochondria with inclusion bodies in the distal and proximal dendrites of VApc and CM in control and parkinsonian monkeys (J-M). Scale bars: A–H = 2μm. In the 3D images, dendrites are shown in white, whereas degenerated mitochondria are shown in black.

## Discussion

In this study, we used the SBF-SEM 3D electron microscopy approach to quantify and compare the synaptic innervation of thalamocortical neurons by putative corticothalamic (RS terminals), pallidothalamic (F3 terminals) and reticular (F1 terminals) axon terminals in the basal ganglia-receiving regions of the VApc and CM between control and MPTP-treated parkinsonian monkeys. Four main conclusions can be drawn from our observations. First, the general pattern of innervation of VApc and CM neurons by putative cortical, pallidal and reticular neurons is not altered in MPTP-treated parkinsonian monkeys. Second, the relative density of glutamatergic cortical terminals (RS) in contact with distal dendrites is significantly higher in VApc than in CM in both control and parkinsonian monkeys. Third, the volume of putative corticothalamic terminals is significantly increased in the VApc and CM of parkinsonian monkeys. Fourth, the complexity and ultrastructural integrity of dendritic mitochondria is altered in CM, but not in the VApc, of parkinsonian monkeys. Together with results of our recent studies (Swain et al., 2020; Masilamoni et al., 2024), these findings provide additional evidence that the synaptic innervation of thalamocortical neurons undergoes neuroplastic changes that may contribute to functional dysregulation of the motor basal ganglia-thalamocortical system in the parkinsonian state.

### Glutamatergic innervation of VApc and CM in control Monkeys

The general distribution of RS glutamatergic and F1/F3 GABAergic terminals described in our study is consistent with previous findings from our laboratory and others in rats, cats and monkeys (Grofova and Rinvik, 1974; Kultas-Ilinsky et al., 1983; Smith et al., 1987; Balercia et al., 1996; Sidibe et al., 1997; Ilinsky et al., 1999; Kakei et al., 2001; Jones, 2007; Rovo et al., 2012; Swain et al., 2020). In line with these findings, our quantitative 3D EM analysis demonstrates that the RS (putative layer VI corticothalamic) terminals are ∼ 6 times more abundant than GABAergic RTN (F1) and GPi (F3) terminals on the distal dendrites of VApc projection neurons.

The ultrastructural features and potential sources of RS terminals have been characterized in detail by various groups using a range of tract tracing approaches, immunohistochemistry and electron microscopy (Jones, 2007; Acsady, 2024). The strong prevalence of RS corticothalamic boutons in VApc is consistent with neuroanatomical data showing that the basal ganglia-receiving nuclei of the primate motor thalamus receive massive glutamatergic inputs from layer VI neurons in the primary motor cortex, the supplementary motor area, and the premotor cortex (Rascol et al., 1992; Rouiller et al., 1998; Rouiller and Welker, 2000; McFarland and Haber, 2002). Another potential source of cortical terminals to the VApc could be axon collaterals of layer V corticospinal neurons. Single cell filling and retrograde viral transduction experiments have, indeed, demonstrated that pyramidal tract (PT) corticospinal axons from M1 send profuse collateral projections to the motor and sensory thalamus in rodents (Kita and Kita, 2012; Sinopoulou et al., 2022). However, recent evidence showed that PT axon collaterals innervation of the motor thalamus in primates is far less prominent than in rodents, suggesting important species differences in the cellular sources of cortical inputs to the ventral motor thalamus (Sinopoulou et al., 2022). Considering studies from other thalamic regions that receive prominent inputs from layer V corticofugal axon collaterals, there is consensus that these terminals are much larger and form many more synapses than RS terminals examined in our study. Significant differences in synaptic strength and release properties between these two populations of cortical terminals have also been demonstrated; the RS terminals being synaptically weak and considered as modulators, while terminals from layer V neurons are synaptically strong and seen as thalamic drivers (Bourassa et al., 1995; Smith and Pare, 1996; Sherman and Guillery, 1998; Rovo et al. 2012; Acsady, 2024). Given evidence that the ventral motor thalamus is devoid of large “driver-like” cortical terminals (Rovo et al., 2012), it is likely that the main source of cortical inputs to the VApc are the layer VI RS corticothalamic neurons. Alternatively, it is possible that some of the terminals categorized as RS in our study originate from layer V axon collaterals, but do not display the typical driver-like multisynaptic ultrastructure in the primate basal ganglia-receiving VApc. Future studies using tract-tracing and physiological methods that allow differentiating layer V from layer VI corticothalamic projections to the VApc are needed to further examine the specific properties of these cortical inputs in the motor thalamus.

Although the pattern of CM neurons’ innervation by the three populations of terminals examined in this study is consistent with our findings in VApc, one striking difference that stood out from our 3D quantitative reconstruction analysis is the 5-6 fold difference in the density of RS terminals in contact with distal dendrites in CM compared to VApc. Overall, the difference in the extent of distal vs proximal dendritic innervation of CM neurons by RS terminals was far less pronounced in CM than VApc. This observation, which confirms and extends earlier electron microscopy results (Balercia et al.,1996), is consistent with data of tract tracing studies showing that the VApc receives a far stronger innervation than CM from motor cortical areas, except for the lateralmost region of CM which harbors a stronger M1 innervation than other CM sub-regions (Smith and Parent, 1986; Kultas-Ilinsky et al., 2003). Thus, considering that the blocks of tissue collected from CM in our study were in the central core of the nucleus, we cannot rule out that the prevalence of RS terminals in CM presented in this study may be lower than what would have been found if tissue blocks had been taken more laterally. Because of the limited amount of data on the electrophysiological effects of cortical inputs on VApc vs CM neurons, the functional significance of these observations remains to be established.

### Ultrastructural remodeling of corticothalamic terminals in the parkinsonian state

We recently showed that the synaptic architecture of GABAergic pallidothalamic terminals in the motor thalamus of parkinsonian monkeys undergo complex ultrastructural changes that may contribute to the increased tonic inhibitory outflow of GPi upon thalamocortical neurons in the parkinsonian state (Masilamoni et al., 2024). The results of the present study suggest that corticothalamic terminals in both VApc and CM also undergo morphological neuroplastic changes in the parkinsonian state. 3D EM reconstruction of over 1300 RS terminals revealed a significant increase in volume of these boutons in parkinsonian monkeys. These findings extend and strengthen our recent single section-based data suggesting an increased cross-sectional diameter of vGluT1-positive corticothalamic terminals in VApc and CM of parkinsonian monkeys (Swain et al. 2019). Combined with our recent data suggesting a change in the density of corticothalamic terminals in the VApc of MPTP-treated parkinsonian monkeys (Swain et al., 2020), these observations suggest that the corticothalamic system undergoes significant neuroplasticity in parkinsonism. Given the limited information on the properties of corticothalamic synapses in VApc and CM in the control parkinsonian state, the functional significance of an increased RS terminal volume remains speculative. However, one might suggest that such a structural change may allow RS corticothalamic terminals to store and possibly release larger amount of glutamate upon increased firing rate. It may also provide terminals the machinery to sustain longer and stronger period of synaptic activation without synaptic vesicles depletion, thereby allowing for increased neuromodulatory influences of RS terminals upon VApc and CM neurons. Although the size of asymmetric synapses formed by RS terminals was not measured in this study, it is possible that the increased capacity of glutamate release and storage in enlarged RS terminals is accompanied with an expansion of their postsynaptic density, an additional structural feature associated with increased synaptic strength. Whether these morphological changes are the primary source of corticothalamic dysfunction or homeostatic responses of RS terminals to other physiological changes of VApc and CM neurons activity is unclear. In light of recent evidence for a robust breakdown of the thalamic innervation of motor cortices in rodent and nonhuman primate models of PD (Villalba et al., 2021; Chen et al., 2023), these results indicate that the thalamo-cortico-thalamic motor circuit is anatomically disrupted in parkinsonism. Considering the complex changes in firing rate, firing pattern and oscillatory activity ventral motor and CM thalamic neurons undergo in PD and animals models of parkinsonism (Schneider and Rothblat, 1996; Molnar et al., 2005; Pessiglione et al., 2005; Bosch-Bouju et al., 2014; Wang et al., 2016; Singh, 2018; Li et al., 2022), future studies that bridge structure-function of synaptic afferents to changes in firing activity could help further determine the underlying synaptic dysregulation involved in the pathophysiology of the thalamo-cortico-thalamic motor loop in PD.

### Dendritic mitochondrial morphology alterations in the VApc and CM of parkinsonian monkeys

Given evidence that dendritic mitochondria are critically important for energy supply, plasticity and proper function of synapses, Ca^2+^ regulation, dendritic formation and other major signaling mechanisms (Verstreken et al., 2005; Ivannikov et al., 2013; Sun et al., 2013; Smith et al., 2016).(Attwell and Laughlin, 2001; Howarth et al., 2012) (Augustine et al., 2003; Grienberger and Konnerth, 2012). (Li et al., 2004; Chang et al., 2006; Ishihara et al., 2009) (Ishihara et al., 2009; Kimura and Murakami, 2014; Fukumitsu et al., 2015), we characterized the morphology, prevalence and pathology of mitochondria in dendrites of VApc and CM neurons in control and parkinsonian monkeys. In contrast to single EM section analysis which offers a partial view of mitochondria profiles, the 3D EM reconstruction approach used in our study allowed for a detailed morphometric analysis of fully reconstructed mitochondria. Overall, our data showed a major difference in the number, complexity and pathology of dendritic mitochondria between control and parkinsonian monkeys in CM, but not in VApc such that distal dendrites of CM neurons in parkinsonian animals underwent a significant numerical decrease, reduced complexity and increased number of mitochondria with dense inclusion bodies, while no difference was found in VApc. Although the functional significance of these observations remains to be elucidated, potential changes in energy supply and metabolic function of CM neurons in the MPTP-induced parkinsonian state should be considered. Interestingly, CM neurons undergo profound degeneration in PD (Henderson et al., 2000; Halliday et al., 2005; Halliday, 2009; Villalba et al., 2014), which is not the case for VApc and most thalamic nuclei. It has also been shown that the loss of CM neurons in PD is unlikely to be induced by increased synuclein pathology, thereby indicating that other cell death mechanisms are involved (Halliday et al., 2005; Halliday, 2009). Combined with our previous findings showing CM neuronal degeneration in chronically MPTP-treated monkeys (Villalba et al., 2014), one may speculate that a mitochondrial energy breakdown might contribute to the death of CM neurons in PD.

### Concluding Remarks

Findings reported in this study further demonstrate that the basal-ganglia-receiving regions of the motor thalamus undergo various forms of synaptic plasticity and, in the case of CM, mitochondrial pathology, in MPTP-treated parkinsonian monkeys. Considering our limited understanding of thalamic and cortical changes associated with the development of parkinsonism, these data lay the foundation for future functional studies of changes in cortical regulation of motor thalamic neurons in PD. They also pave the way for future analyses of the cellular mechanisms that may contribute to energy supply deficiency, and potentially death, of CM neurons in PD.

## Acknowledgements

This work was supported by the NIH grant P50NS123103 and the NIH/OD/ORIP base grant of the Emory National Primate Research Center P51OD011132. Thanks to Susan Jenkins for technical assistance with tissue processing. Thanks are also due to Emily Benson of Renovo Neural Inc, and now Lerner Research Institute’s 3DEM Core (Cleveland Clinic) for sample preparation and generating SBF/SEM datasets.

---

**Supp. Figure 1.** Photomicrographs of TH-immunostained coronal sections at the level of the pre-commissural striatum (A–B), post-commissural striatum (C–D), and midbrain dopaminergic cell groups (E–F) of a control (left column) and a MPTP-treated (right column) monkey. Note the profound loss of TH innervation in the post-commissural striatum and ventral tier of the SNc. Abbreviations: CA: caudate nucleus; GPe: globus pallidus, external segment; GPi: globus pallidus, internal segment; LC: locus coeruleus; PU: putamen; SNCd: substantia nigra compacta, dorsal tier; SNCv: substantia nigra, ventral tier; Th: thalamus; VTA: ventral tegmental area. Scale bars: A–D = 5mm, E-F = 2mm.

**Supp. Figure 2.** Representative FIB/SEM images showing the three main types of axon terminals contacting neuronal cell bodies. (A) RS-type terminal, (B) F1-type terminal, and (C) F3-type terminal. Scatter dot plots with bar graphs comparing the total terminal densities of RS, F1, and F3 terminals in the cell body of the VApc (D) and CM (E) between control and MPTP-treated parkinsonian monkeys. No total terminal density differences were found for the three types of terminals between control and parkinsonian monkeys.

## Statistical Table

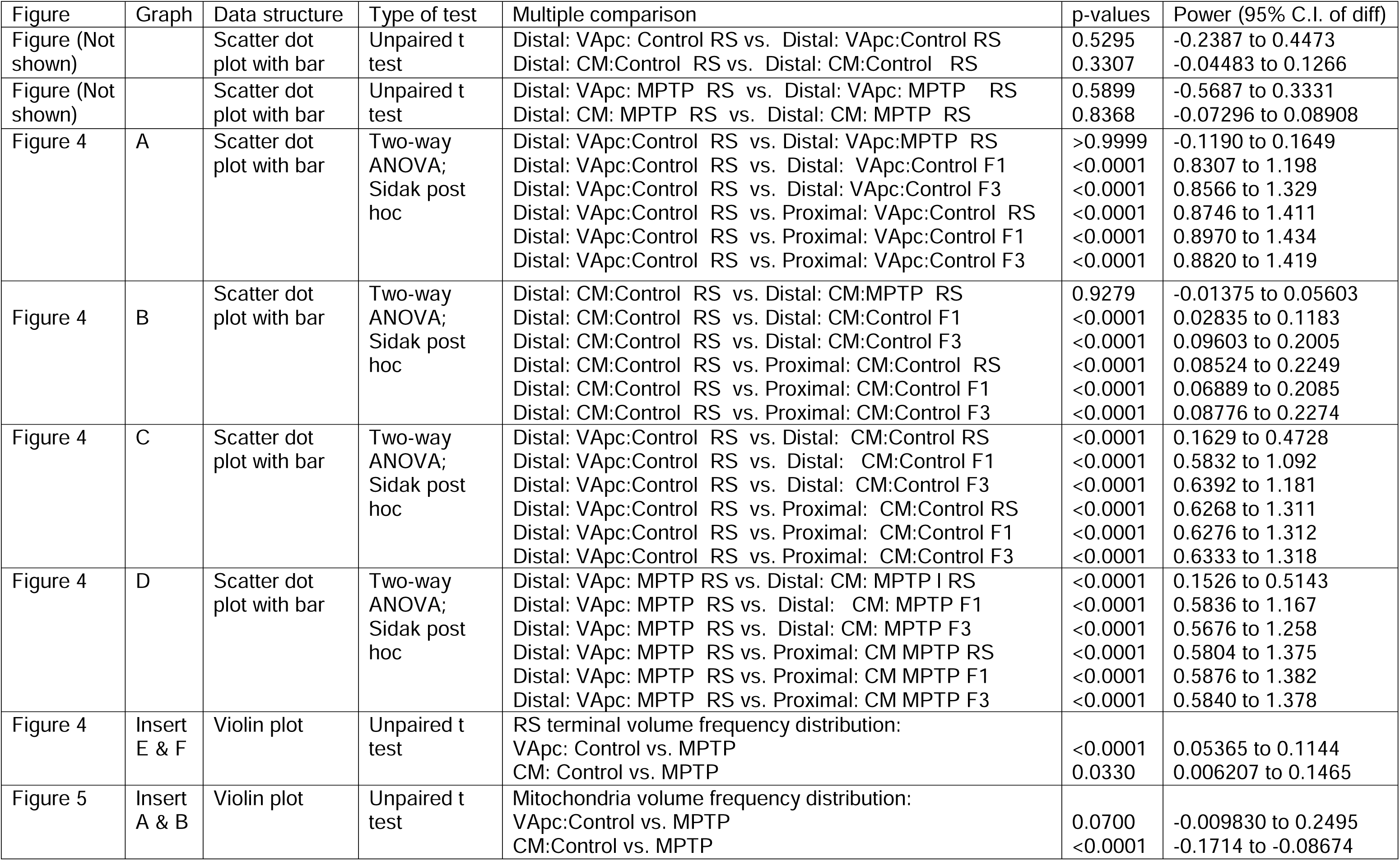
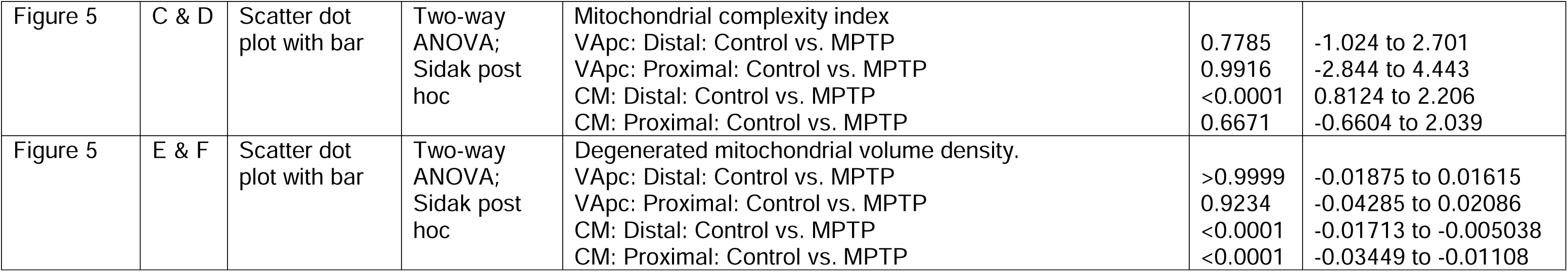

| Figure | Graph | Data structure | Type of test | Multiple comparison | p-values | Power (95% C.I. of diff) |
| --- | --- | --- | --- | --- | --- | --- |
| Figure (Not shown) |  | Scatter dot plot with bar | Unpaired t test | Distal: VApc: Control RS vs. Distal: VApc:Control RS<br>Distal: CM:Control RS vs. Distal: CM:Control RS | 0.5295<br>0.3307 | -0.2387 to 0.4473<br>-0.04483 to 0.1266 |
| Figure (Not shown) |  | Scatter dot plot with bar | Unpaired t test | Distal: VApc: MPTP RS vs. Distal: VApc: MPTP RS<br>Distal: CM: MPTP RS vs. Distal: CM: MPTP RS | 0.5899<br>0.8368 | -0.5687 to 0.3331<br>-0.07296 to 0.08908 |
| Figure 4 | A | Scatter dot plot with bar | Two-way ANOVA;<br>Sidak post hoc | Distal: VApc:Control RS vs. Distal: VApc:MPTP RS<br>Distal: VApc:Control RS vs. Distal: VApc:Control F1<br>Distal: VApc:Control RS vs. Distal: VApc:Control F3<br>Distal: VApc:Control RS vs. Proximal: VApc:Control RS<br>Distal: VApc:Control RS vs. Proximal: VApc:Control F1<br>Distal: VApc:Control RS vs. Proximal: VApc:Control F3 | >0.9999<br><0.0001<br><0.0001<br><0.0001<br><0.0001<br><0.0001 | -0.1190 to 0.1649<br>0.8307 to 1.198<br>0.8566 to 1.329<br>0.8746 to 1.411<br>0.8970 to 1.434<br>0.8820 to 1.419 |
| Figure 4 | B | Scatter dot plot with bar | Two-way ANOVA;<br>Sidak post hoc | Distal: CM:Control RS vs. Distal: CM:MPTP RS<br>Distal: CM:Control RS vs. Distal: CM:Control F1<br>Distal: CM:Control RS vs. Distal: CM:Control F3<br>Distal: CM:Control RS vs. Proximal: CM:Control RS<br>Distal: CM:Control RS vs. Proximal: CM:Control F1<br>Distal: CM:Control RS vs. Proximal: CM:Control F3 | 0.9279<br><0.0001<br><0.0001<br><0.0001<br><0.0001<br><0.0001 | -0.01375 to 0.05603<br>0.02835 to 0.1183<br>0.09603 to 0.2005<br>0.08524 to 0.2249<br>0.06889 to 0.2085<br>0.08776 to 0.2274 |
| Figure 4 | C | Scatter dot plot with bar | Two-way ANOVA;<br>Sidak post hoc | Distal: VApc:Control RS vs. Distal: CM:Control RS<br>Distal: VApc:Control RS vs. Distal: CM:Control F1<br>Distal: VApc:Control RS vs. Distal: CM:Control F3<br>Distal: VApc:Control RS vs. Proximal: CM:Control RS<br>Distal: VApc:Control RS vs. Proximal: CM:Control F1<br>Distal: VApc:Control RS vs. Proximal: CM:Control F3 | <0.0001<br><0.0001<br><0.0001<br><0.0001<br><0.0001<br><0.0001 | 0.1629 to 0.4728<br>0.5832 to 1.092<br>0.6392 to 1.181<br>0.6268 to 1.311<br>0.6276 to 1.312<br>0.6333 to 1.318 |
| Figure 4 | D | Scatter dot plot with bar | Two-way ANOVA;<br>Sidak post hoc | Distal: VApc: MPTP RS vs. Distal: CM: MPTP I RS<br>Distal: VApc: MPTP RS vs. Distal: CM: MPTP F1<br>Distal: VApc: MPTP RS vs. Distal: CM: MPTP F3<br>Distal: VApc: MPTP RS vs. Proximal: CM MPTP RS<br>Distal: VApc: MPTP RS vs. Proximal: CM MPTP F1<br>Distal: VApc: MPTP RS vs. Proximal: CM MPTP F3 | <0.0001<br><0.0001<br><0.0001<br><0.0001<br><0.0001<br><0.0001 | 0.1526 to 0.5143<br>0.5836 to 1.167<br>0.5676 to 1.258<br>0.5804 to 1.375<br>0.5876 to 1.382<br>0.5840 to 1.378 |
| Figure 4 | Insert E & F | Violin plot | Unpaired t test | RS terminal volume frequency distribution:<br>VApc: Control vs. MPTP<br>CM: Control vs. MPTP | <0.0001<br>0.0330 | 0.05365 to 0.1144<br>0.006207 to 0.1465 |
| Figure 5 | Insert A & B | Violin plot | Unpaired t test | Mitochondria volume frequency distribution:<br>VApc:Control vs. MPTP<br>CM:Control vs. MPTP | 0.0700<br><0.0001 | -0.009830 to 0.2495<br>-0.1714 to -0.08674 |
| Statistical Table |  |  |  |  |  |  |
| Figure 5 | C & D | Scatter dot plot with bar | Two-way ANOVA; Sidak post hoc | Mitochondrial complexity index<br>VApc: Distal: Control vs. MPTP<br>VApc: Proximal: Control vs. MPTP<br>CM: Distal: Control vs. MPTP<br>CM: Proximal: Control vs. MPTP | 0.7785<br>0.9916<br><0.0001<br>0.6671 | -1.024 to 2.701<br>-2.844 to 4.443<br>0.8124 to 2.206<br>-0.6604 to 2.039 |
| Figure 5 | E & F | Scatter dot plot with bar | Two-way ANOVA; Sidak post hoc | Degenerated mitochondrial volume density.<br>VApc: Distal: Control vs. MPTP<br>VApc: Proximal: Control vs. MPTP<br>CM: Distal: Control vs. MPTP<br>CM: Proximal: Control vs. MPTP | >0.9999<br>0.9234<br><0.0001<br><0.0001 | -0.01875 to 0.01615<br>-0.04285 to 0.02086<br>-0.01713 to -0.005038<br>-0.03449 to -0.01108 |

## REFERENCES

1. Adoff MD, Climer JR, Davoudi H, Marvin JS, Looger LL, Dombeck DA (2021) The functional organization of excitatory synaptic input to place cells. Nat Commun 12:3558.

2. Attwell D, Laughlin SB (2001) An energy budget for signaling in the grey matter of the brain. J Cereb Blood Flow Metab 21:1133–1145.

3. Augustine GJ, Santamaria F, Tanaka K (2003) Local calcium signaling in neurons. Neuron 40:331–346.

4. Aymerich MS, Barroso-Chinea P, Perez-Manso M, Munoz-Patino AM, Moreno-Igoa M, Gonzalez-Hernandez T, Lanciego JL (2006) Consequences of unilateral nigrostriatal denervation on the thalamostriatal pathway in rats. Eur J Neurosci 23:2099–2108.

5. Balercia G, Kultas-Ilinsky K, Bentivoglio M, Ilinsky IA (1996) Neuronal and synaptic organization of the centromedian nucleus of the monkey thalamus: a quantitative ultrastructural study, with tract tracing and immunohistochemical observations. J Neurocytol 25:267–288.

6. Bosch-Bouju C, Hyland BI, Parr-Brownlie LC (2013) Motor thalamus integration of cortical, cerebellar and basal ganglia information: implications for normal and parkinsonian conditions. Front Comput Neurosci 7:163.

7. Bosch-Bouju C, Smither RA, Hyland BI, Parr-Brownlie LC (2014) Reduced reach-related modulation of motor thalamus neural activity in a rat model of Parkinson’s disease. J Neurosci 34:15836–15850.

8. Chang DT, Honick AS, Reynolds IJ (2006) Mitochondrial trafficking to synapses in cultured primary cortical neurons. J Neurosci 26:7035–7045.

9. Chen H, Zhuang P, Miao SH, Yuan G, Zhang YQ, Li JY, Li YJ (2010) Neuronal firing in the ventrolateral thalamus of patients with Parkinson’s disease differs from that with essential tremor. Chin Med J (Engl) 123:695–701.

10. Cheng A, Hou Y, Mattson MP (2010) Mitochondria and neuroplasticity. ASN Neuro 2:e00045.

11. Christie-Pope BC, Burns RS, Whetsell WO, Jr. (1989) Ultrastructural alterations induced by 1-methyl-4-phenylpyridinium (MPP+) in canine substantia nigra and rat mesencephalon in vitro. Exp Neurol 104:235–240.

12. Chu HY (2020) Synaptic and cellular plasticity in Parkinson’s disease. Acta Pharmacol Sin 41:447–452.

13. Chu HY, McIver EL, Kovaleski RF, Atherton JF, Bevan MD (2017) Loss of Hyperdirect Pathway Cortico-Subthalamic Inputs Following Degeneration of Midbrain Dopamine Neurons. Neuron 95:1306–1318 e1305.

14. Day M, Wang Z, Ding J, An X, Ingham CA, Shering AF, Wokosin D, Ilijic E, Sun Z, Sampson AR, Mugnaini E, Deutch AY, Sesack SR, Arbuthnott GW, Surmeier DJ (2006) Selective elimination of glutamatergic synapses on striatopallidal neurons in Parkinson disease models. Nat Neurosci 9:251–259.

15. Delgado T, Petralia RS, Freeman DW, Sedlacek M, Wang YX, Brenowitz SD, Sheu SH, Gu JW, Kapogiannis D, Mattson MP, Yao PJ (2019) Comparing 3D ultrastructure of presynaptic and postsynaptic mitochondria. Biol Open 8.

16. DeLong MR (1990) Primate models of movement disorders of basal ganglia origin. Trends Neurosci 13:281–285.

17. Duchen MR (2000) Mitochondria and Ca(2+)in cell physiology and pathophysiology. Cell Calcium 28:339–348.

18. Faitg J, Lacefield C, Davey T, White K, Laws R, Kosmidis S, Reeve AK, Kandel ER, Vincent AE, Picard M (2021) 3D neuronal mitochondrial morphology in axons, dendrites, and somata of the aging mouse hippocampus. Cell Rep 36:109509.

19. Fukumitsu K, Fujishima K, Yoshimura A, Wu YK, Heuser J, Kengaku M (2015) Synergistic action of dendritic mitochondria and creatine kinase maintains ATP homeostasis and actin dynamics in growing neuronal dendrites. J Neurosci 35:5707–5723.

20. Grienberger C, Konnerth A (2012) Imaging calcium in neurons. Neuron 73:862–885.

21. Grofova I, Rinvik E (1974) Cortical and pallidal projections to the nucleus ventralis lateralis thalami. Electron microscopical studies in the cat. Anat Embryol (Berl) 146:113–132.

22. Guehl D, Pessiglione M, Francois C, Yelnik J, Hirsch EC, Feger J, Tremblay L (2003) Tremor-related activity of neurons in the ’motor’ thalamus: changes in firing rate and pattern in the MPTP vervet model of parkinsonism. Eur J Neurosci 17:2388–2400.

23. Halliday GM (2009) Thalamic changes in Parkinson’s disease. Parkinsonism Relat Disord 15 Suppl 3:S152–155.

24. Halliday GM, Macdonald V, Henderson JM (2005) A comparison of degeneration in motor thalamus and cortex between progressive supranuclear palsy and Parkinson’s disease. Brain 128:2272–2280.

25. Hamori J, Pasik T, Pasik P, Szentagothai J (1974) Triadic synaptic arrangements and their possible significance in the lateral geniculate nucleus of the monkey. Brain Res 80:379–393.

26. Hayes MJ, Tracey-White D, Kam JH, Powner MB, Jeffery G (2021) The 3D organisation of mitochondria in primate photoreceptors. Sci Rep 11:18863.

27. Henderson JM, Carpenter K, Cartwright H, Halliday GM (2000) Degeneration of the centre median-parafascicular complex in Parkinson’s disease. Ann Neurol 47:345–352.

28. Howarth C, Gleeson P, Attwell D (2012) Updated energy budgets for neural computation in the neocortex and cerebellum. J Cereb Blood Flow Metab 32:1222–1232.

29. Iascone DM, Li Y, Sumbul U, Doron M, Chen H, Andreu V, Goudy F, Blockus H, Abbott LF, Segev I, Peng H, Polleux F (2020) Whole-Neuron Synaptic Mapping Reveals Spatially Precise Excitatory/Inhibitory Balance Limiting Dendritic and Somatic Spiking. Neuron 106:566–578 e568.

30. Ilinsky IA, Yi H, Kultas-Ilinsky K (1997) Mode of termination of pallidal afferents to the thalamus: a light and electron microscopic study with anterograde tracers and immunocytochemistry in Macaca mulatta. J Comp Neurol 386:601–612.

31. Ilinsky IA, Ambardekar AV, Kultas-Ilinsky K (1999) Organization of projections from the anterior pole of the nucleus reticularis thalami (NRT) to subdivisions of the motor thalamus: light and electron microscopic studies in the rhesus monkey. J Comp Neurol 409:369–384.

32. Ingham CA, Hood SH, Taggart P, Arbuthnott GW (1998) Plasticity of synapses in the rat neostriatum after unilateral lesion of the nigrostriatal dopaminergic pathway. J Neurosci 18:4732–4743.

33. Ishihara N, Nomura M, Jofuku A, Kato H, Suzuki SO, Masuda K, Otera H, Nakanishi Y, Nonaka I, Goto Y, Taguchi N, Morinaga H, Maeda M, Takayanagi R, Yokota S, Mihara K (2009) Mitochondrial fission factor Drp1 is essential for embryonic development and synapse formation in mice. Nat Cell Biol 11:958–966.

34. Ivannikov MV, Sugimori M, Llinas RR (2013) Synaptic vesicle exocytosis in hippocampal synaptosomes correlates directly with total mitochondrial volume. J Mol Neurosci 49:223–230.

35. Ji YW, Zhang X, Fan JP, Gu WX, Shen ZL, Wu HC, Cui G, Zhou C, Xiao C (2023) Differential remodeling of subthalamic projections to basal ganglia output nuclei and locomotor deficits in 6-OHDA-induced hemiparkinsonian mice. Cell Rep 42:112178.

36. Johnson MD, Thompson CK, Tysseling VM, Powers RK, Heckman CJ (2017) The potential for understanding the synaptic organization of human motor commands via the firing patterns of motoneurons. J Neurophysiol 118:520–531.

37. Jones E (2007) The Thalamus (2nd edn.). Cambridge (UK): Cambridge University Press.

38. Jones EG (2002) Thalamic organization and function after Cajal. Prog Brain Res 136:333–357.

39. Kakei S, Na J, Shinoda Y (2001) Thalamic terminal morphology and distribution of single corticothalamic axons originating from layers 5 and 6 of the cat motor cortex. J Comp Neurol 437:170–185.

40. Kammermeier S, Pittard D, Hamada I, Wichmann T (2016) Effects of high-frequency stimulation of the internal pallidal segment on neuronal activity in the thalamus in parkinsonian monkeys. J Neurophysiol 116:2869–2881.

41. Kimura T, Murakami F (2014) Evidence that dendritic mitochondria negatively regulate dendritic branching in pyramidal neurons in the neocortex. J Neurosci 34:6938–6951.

42. Kultas-Ilinsky K, Ilinsky I, Warton S, Smith KR (1983) Fine structure of nigral and pallidal afferents in the thalamus: an EM autoradiography study in the cat. J Comp Neurol 216:390–405.

43. Kultas-Ilinsky K, Reising L, Yi H, Ilinsky IA (1997) Pallidal afferent territory of the Macaca mulatta thalamus: neuronal and synaptic organization of the VAdc. J Comp Neurol 386:573–600.

44. Kuzniewska B, Cysewski D, Wasilewski M, Sakowska P, Milek J, Kulinski TM, Winiarski M, Kozielewicz P, Knapska E, Dadlez M, Chacinska A, Dziembowski A, Dziembowska M (2020) Mitochondrial protein biogenesis in the synapse is supported by local translation. EMBO Rep 21:e48882.

45. Li H, Li SH, Yu ZX, Shelbourne P, Li XJ (2001) Huntingtin aggregate-associated axonal degeneration is an early pathological event in Huntington’s disease mice. J Neurosci 21:8473–8481.

46. Li M, Zhang X, He Q, Chen D, Chen F, Wang X, Sun S, Sun Y, Li Y, Zhu Z, Fang H, Shi X, Yao X, Sun H, Wang M (2022) Functional Interactions Between the Parafascicular Thalamic Nucleus and Motor Cortex Are Altered in Hemiparkinsonian Rat. Front Aging Neurosci 14:800159.

47. Li Z, Okamoto K, Hayashi Y, Sheng M (2004) The importance of dendritic mitochondria in the morphogenesis and plasticity of spines and synapses. Cell 119:873–887.

48. Magnin M, Morel A, Jeanmonod D (2000) Single-unit analysis of the pallidum, thalamus and subthalamic nucleus in parkinsonian patients. Neuroscience 96:549–564.

49. Masilamoni G, Kelly H, Swain A, Y S (2021) Ultrastructural plasticity of pallidothalamic terminals in the motor thalamus of MPTP-treated parkinsonian monkeys: A 3D quantitative analysis. In: Soc. Neurosci. Abstr. .

50. Masilamoni GJ, Kelly H, Swain AJ, Pare JF, Villalba RM, Smith Y (2024) Structural Plasticity of GABAergic Pallidothalamic Terminals in MPTP-Treated Parkinsonian Monkeys: A 3D Electron Microscopic Analysis. eNeuro 11.

51. Mathai A, Smith Y (2011) The corticostriatal and corticosubthalamic pathways: two entries, one target. So what? Front Syst Neurosci 5:64.

52. Mathai A, Ma Y, Pare JF, Villalba RM, Wichmann T, Smith Y (2015) Reduced cortical innervation of the subthalamic nucleus in MPTP-treated parkinsonian monkeys. Brain 138:946–962.

53. McFarland NR, Haber SN (2002) Thalamic relay nuclei of the basal ganglia form both reciprocal and nonreciprocal cortical connections, linking multiple frontal cortical areas. J Neurosci 22:8117–8132.

54. Melief EJ, McKinley JW, Lam JY, Whiteley NM, Gibson AW, Neumaier JF, Henschen CW, Palmiter RD, Bamford NS, Darvas M (2018) Loss of glutamate signaling from the thalamus to dorsal striatum impairs motor function and slows the execution of learned behaviors. NPJ Parkinsons Dis 4:23.

55. Merchan-Perez A, Rodriguez JR, Alonso-Nanclares L, Schertel A, Defelipe J (2009) Counting Synapses Using FIB/SEM Microscopy: A True Revolution for Ultrastructural Volume Reconstruction. Front Neuroanat 3:18.

56. Merino-Galan L, Jimenez-Urbieta H, Zamarbide M, Rodriguez-Chinchilla T, Belloso-Iguerategui A, Santamaria E, Fernandez-Irigoyen J, Aiastui A, Doudnikoff E, Bezard E, Ouro A, Knafo S, Gago B, Quiroga-Varela A, Rodriguez-Oroz MC (2022) Striatal synaptic bioenergetic and autophagic decline in premotor experimental parkinsonism. Brain 145:2092–2107.

57. Mizukawa K, Sora YH, Ogawa N (1990) Ultrastructural changes of the substantia nigra, ventral tegmental area and striatum in 1-methyl-4-phenyl-1,2,3,6-tetrahydropyridine (MPTP)-treated mice. Res Commun Chem Pathol Pharmacol 67:307–320.

58. Molnar GF, Pilliar A, Lozano AM, Dostrovsky JO (2005) Differences in neuronal firing rates in pallidal and cerebellar receiving areas of thalamus in patients with Parkinson’s disease, essential tremor, and pain. J Neurophysiol 93:3094–3101.

59. Montero T, Gatica RI, Farassat N, Meza R, Gonzalez-Cabrera C, Roeper J, Henny P (2021) Dendritic Architecture Predicts in vivo Firing Pattern in Mouse Ventral Tegmental Area and Substantia Nigra Dopaminergic Neurons. Front Neural Circuits 15:769342.

60. Ni ZG, Gao DM, Benabid AL, Benazzouz A (2000) Unilateral lesion of the nigrostriatal pathway induces a transient decrease of firing rate with no change in the firing pattern of neurons of the parafascicular nucleus in the rat. Neuroscience 101:993–999.

61. Nicholls DG, Budd SL (2000) Mitochondria and neuronal survival. Physiol Rev 80:315–360.

62. Obeso JA, Olanow CW, Nutt JG (2000) Levodopa motor complications in Parkinson’s disease. Trends Neurosci 23:S2–7.

63. Pan K, Jinnah HA, Hess EJ, Smith Y, Villalba RM (2023) Ultrastructural analysis of nigrostriatal dopaminergic terminals in a knockin mouse model of DYT1 dystonia. Eur J Neurosci:e16197.

64. Pessiglione M, Guehl D, Rolland AS, Francois C, Hirsch EC, Feger J, Tremblay L (2005) Thalamic neuronal activity in dopamine-depleted primates: evidence for a loss of functional segregation within basal ganglia circuits. J Neurosci 25:1523–1531.

65. Peters A, Palay SL, Webster de F (1991) The Fine Structure of the Nervous System: Neurons and Their Supporting Cells. In.

66. Raeva S, Vainberg N, Dubinin V (1999) Analysis of spontaneous activity patterns of human thalamic ventrolateral neurons and their modifications due to functional brain changes. Neuroscience 88:365–376.

67. Raju DV, Ahern TH, Shah DJ, Wright TM, Standaert DG, Hall RA, Smith Y (2008) Differential synaptic plasticity of the corticostriatal and thalamostriatal systems in an MPTP-treated monkey model of parkinsonism. Eur J Neurosci 27:1647–1658.

68. Rangaraju V, Lauterbach M, Schuman EM (2019) Spatially Stable Mitochondrial Compartments Fuel Local Translation during Plasticity. Cell 176:73–84 e15.

69. Rapisardi SC, Warrington VO, Wilson JS (1990) Effects of MPTP on the fine structure of neurons in substantia nigra of dogs. Brain Res 512:147–154.

70. Rascol O, Sabatini U, Chollet F, Celsis P, Montastruc JL, Marc-Vergnes JP, Rascol A (1992) Supplementary and primary sensory motor area activity in Parkinson’s disease. Regional cerebral blood flow changes during finger movements and effects of apomorphine. Arch Neurol 49:144–148.

71. Rolland AS, Herrero MT, Garcia-Martinez V, Ruberg M, Hirsch EC, Francois C (2007) Metabolic activity of cerebellar and basal ganglia-thalamic neurons is reduced in parkinsonism. Brain 130:265–275.

72. Rouiller EM, Welker E (2000) A comparative analysis of the morphology of corticothalamic projections in mammals. Brain Res Bull 53:727–741.

73. Rouiller EM, Tanne J, Moret V, Kermadi I, Boussaoud D, Welker E (1998) Dual morphology and topography of the corticothalamic terminals originating from the primary, supplementary motor, and dorsal premotor cortical areas in macaque monkeys. J Comp Neurol 396:169–185.

74. Rovo Z, Ulbert I, Acsady L (2012) Drivers of the primate thalamus. J Neurosci 32:17894–17908.

75. Rubin JE, McIntyre CC, Turner RS, Wichmann T (2012) Basal ganglia activity patterns in parkinsonism and computational modeling of their downstream effects. Eur J Neurosci 36:2213–2228.

76. Ruggiero A, Katsenelson M, Slutsky I (2021) Mitochondria: new players in homeostatic regulation of firing rate set points. Trends Neurosci 44:605–618.

77. Sarnthein J, Jeanmonod D (2007) High thalamocortical theta coherence in patients with Parkinson’s disease. J Neurosci 27:124–131.

78. Scheff SW, Price DA, Schmitt FA, DeKosky ST, Mufson EJ (2007) Synaptic alterations in CA1 in mild Alzheimer disease and mild cognitive impairment. Neurology 68:1501–1508.

79. Schneider JS, Rothblat DS (1996) Alterations in intralaminar and motor thalamic physiology following nigrostriatal dopamine depletion. Brain Res 742:25–33.

80. Sherman SM (2004) Interneurons and triadic circuitry of the thalamus. Trends Neurosci 27:670–675.

81. Sherman SM, Friedlander MJ (1988) Identification of X versus Y properties for interneurons in the A-laminae of the cat’s lateral geniculate nucleus. Exp Brain Res 73:384–392.

82. Sidibe M, Bevan MD, Bolam JP, Smith Y (1997) Efferent connections of the internal globus pallidus in the squirrel monkey: I. Topography and synaptic organization of the pallidothalamic projection. J Comp Neurol 382:323–347.

83. Singh A (2018) Oscillatory activity in the cortico-basal ganglia-thalamic neural circuits in Parkinson’s disease. Eur J Neurosci 48:2869–2878.

84. Smith HL, Bourne JN, Cao G, Chirillo MA, Ostroff LE, Watson DJ, Harris KM (2016) Mitochondrial support of persistent presynaptic vesicle mobilization with age-dependent synaptic growth after LTP. Elife 5.

85. Smith Y, Seguela P, Parent A (1987) Distribution of GABA-immunoreactive neurons in the thalamus of the squirrel monkey (Saimiri sciureus). Neuroscience 22:579–591.

86. Song DD, Shults CW, Sisk A, Rockenstein E, Masliah E (2004) Enhanced substantia nigra mitochondrial pathology in human alpha-synuclein transgenic mice after treatment with MPTP. Exp Neurol 186:158–172.

87. Stoler O, Stavsky A, Khrapunsky Y, Melamed I, Stutzmann G, Gitler D, Sekler I, Fleidervish I (2022) Frequency- and spike-timing-dependent mitochondrial Ca(2+) signaling regulates the metabolic rate and synaptic efficacy in cortical neurons. Elife 11.

88. Sun T, Qiao H, Pan PY, Chen Y, Sheng ZH (2013) Motile axonal mitochondria contribute to the variability of presynaptic strength. Cell Rep 4:413–419.

89. Swain AJ, Galvan A, Wichmann T, Smith Y (2020) Structural plasticity of GABAergic and glutamatergic networks in the motor thalamus of parkinsonian monkeys. J Comp Neurol 528:1436–1456.

90. Tanaka J, Nakamura H, Honda S, Takada K, Kato S (1988) Neuropathological study on 1-methyl-4-phenyl-1,2,3,6-tetrahydropyridine of the crab-eating monkey. Acta Neuropathol 75:370–376.

91. Toescu EC (2000) Mitochondria and Ca(2+) signaling. J Cell Mol Med 4:164–175.

92. Toomey CE, Heywood WE, Evans JR, Lachica J, Pressey SN, Foti SC, Al Shahrani M, D’Sa K, Hargreaves IP, Heales S, Orford M, Troakes C, Attems J, Gelpi E, Palkovits M, Lashley T, Gentleman SM, Revesz T, Mills K, Gandhi S (2022) Mitochondrial dysfunction is a key pathological driver of early stage Parkinson’s. Acta Neuropathol Commun 10:134.

93. Verstreken P, Ly CV, Venken KJ, Koh TW, Zhou Y, Bellen HJ (2005) Synaptic mitochondria are critical for mobilization of reserve pool vesicles at Drosophila neuromuscular junctions. Neuron 47:365–378.

94. Villalba RM, Smith Y (2010) Striatal spine plasticity in Parkinson’s disease. Front Neuroanat 4:133.

95. Villalba RM, Smith Y (2011) Differential structural plasticity of corticostriatal and thalamostriatal axo-spinous synapses in MPTP-treated Parkinsonian monkeys. J Comp Neurol 519:989–1005.

96. Villalba RM, Smith Y (2018) Loss and remodeling of striatal dendritic spines in Parkinson’s disease: from homeostasis to maladaptive plasticity? J Neural Transm (Vienna) 125:431–447.

97. Villalba RM, Lee H, Smith Y (2009) Dopaminergic denervation and spine loss in the striatum of MPTP-treated monkeys. Exp Neurol 215:220–227.

98. Villalba RM, Wichmann T, Smith Y (2014) Neuronal loss in the caudal intralaminar thalamic nuclei in a primate model of Parkinson’s disease. Brain Struct Funct 219:381–394.

99. Villalba RM, Mathai A, Smith Y (2015) Morphological changes of glutamatergic synapses in animal models of Parkinson’s disease. Front Neuroanat 9:117.

100. Vincent AE, White K, Davey T, Philips J, Ogden RT, Lawless C, Warren C, Hall MG, Ng YS, Falkous G, Holden T, Deehan D, Taylor RW, Turnbull DM, Picard M (2019) Quantitative 3D Mapping of the Human Skeletal Muscle Mitochondrial Network. Cell Rep 26:996–1009 e1004.

101. Wang M, Qu Q, He T, Li M, Song Z, Chen F, Zhang X, Xie J, Geng X, Yang M, Wang X, Lei C, Hou Y (2016) Distinct temporal spike and local field potential activities in the thalamic parafascicular nucleus of parkinsonian rats during rest and limb movement. Neuroscience 330:57–71.

102. Wang W, Zhao F, Lu Y, Siedlak SL, Fujioka H, Feng H, Perry G, Zhu X (2023) Damaged mitochondria coincide with presynaptic vesicle loss and abnormalities in alzheimer’s disease brain. Acta Neuropathol Commun 11:54.

103. Wichmann T, DeLong MR (1996) Functional and pathophysiological models of the basal ganglia. Curr Opin Neurobiol 6:751–758.

104. Xu T, Wang S, Lalchandani RR, Ding JB (2017) Motor learning in animal models of Parkinson’s disease: Aberrant synaptic plasticity in the motor cortex. Mov Disord 32:487–497.

105. Zirh TA, Lenz FA, Reich SG, Dougherty PM (1998) Patterns of bursting occurring in thalamic cells during parkinsonian tremor. Neuroscience 83:107–121.

106. Masilamoni, G.J., Kelly, H., Swain, A.J., Pare, J.F., Villalba, R.M. & Smith, Y. (2024) Structural Plasticity of GABAergic Pallidothalamic Terminals in MPTP-Treated Parkinsonian Monkeys: A 3D Electron Microscopic Analysis. eNeuro, 11.

