## Supplementary figures and images for "GABAergic and glutamatergic synaptic networks and mitochondrial morphology in the thalamic ventral motor and centromedian nuclei of Rhesus Monkey: A comparative 3D Electron Microscopic Analysis between Control and Parkinsonian State"

### Suppl figure 1

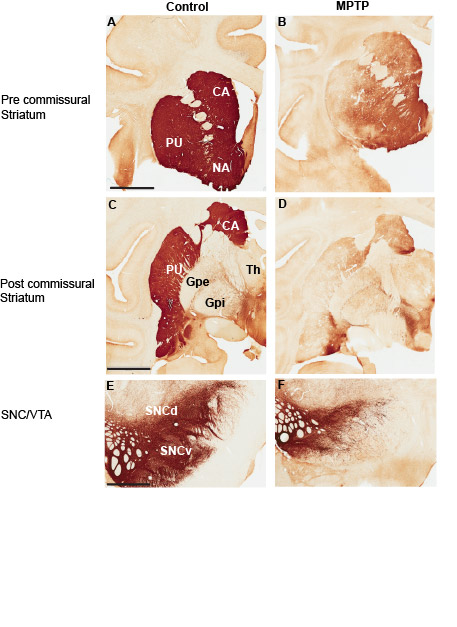

### Suppl figure 2

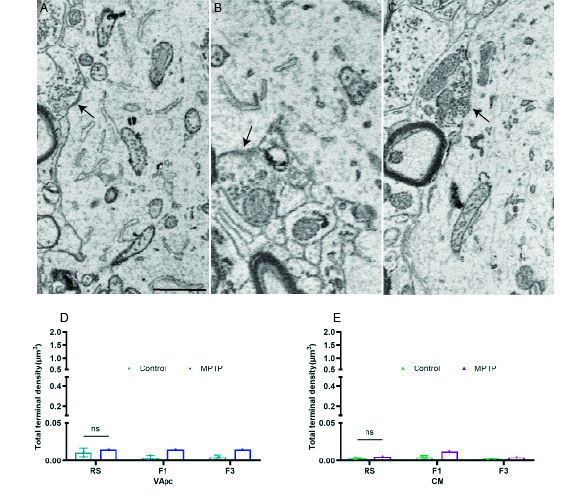
